# Acropetal and basipetal cardenolide transport in *Erysimum cheiranthoides* (wormseed wallflower)

**DOI:** 10.1101/2021.09.03.458893

**Authors:** Martin L. Alani, Gordon C. Younkin, Mahdieh Mirzaei, Pavan Kumar, Georg Jander

## Abstract

Plant specialized metabolites are often subject to within-plant transport and have tissue-specific distribution patterns. Among plants in the Brassicaceae, the genus *Erysimum* is unique in producing not only glucosinolates but also cardenolides as defense against insect herbivory. Ten cardenolides were detected with varying abundance in different tissues of *Erysimum cheiranthoides* (wormseed wallflower). As is predicted by the optimal defense theory, cardenolides were most abundant in young leaves and reproductive tissues. The lowest concentrations were observed in senescing leaves and roots. Crosses between wildtype *E. cheiranthoides* and a mutant line with an altered cardenolide profile showed that the seed cardenolide phenotype is determined entirely by the maternal genotype. Prior to the development of the first true leaves, seedling cotyledons also had the maternal cardenolide profile. Hypocotyl grafting experiments showed that the root cardenolide profile is determined entirely by the aboveground plant genotype. In further grafting experiments, there was no evidence of cardenolide transport into the leaves, but a mixed cardenolide profile was observed in the stems and inflorescences of plants that had been grafted at vegetative and flowering growth stages, respectively. Together, these results indicate that *E. cheiranthoides* leaves are a site of cardenolide biosynthesis and therefore also the plant tissue that is most likely to be expressing the relevant biosynthetic genes.

## 1. Introduction

A wide variety of toxic and deterrent metabolites in leaves, roots, flowers, and other plant parts provide protection against pathogens and herbivores. However, the presence of a defensive metabolite in a particular plant tissue type is not necessarily an indication that it is synthesized there. For instance, nicotine that accumulates in *Nicotiana* spp (tobacco) leaves is produced in the roots (Baldwin, 1989; Cai *et al.,* 2013), glucosinolates are actively transported into developing *Arabidopsis thaliana* seeds (Nour-Eldin *et al.,* 2012), and although colchicine is synthesized in the corms of *Colchicum* spp. (autumn crocus), it is transported throughout the plant (Nett *et al.,* 2020). Given this widespread within-plant movement of defense-related metabolites, any investigation of the biosynthetic pathways of plant specialized metabolism needs to consider the physical location of the biosynthetic enzymes.

Cardiac glycosides are a class of plant defensive metabolites that function as inhibitors of Na^+^, K^+^-ATPase, an essential membrane ion transporter in animal cells (Dzimiri *et al.,* 1987). Biosynthesis of cardiac glycosides, which are broadly classified as cardenolides or bufadienolides based on the presence of a five- or a six-membered lactone ring, evolved independently in at least twelve plant families (Agrawal *et al.,* 2012; Melero *et al.,* 2000; Steyn and van Heerden, 1998). Cardenolides in plants such as *Asclepias* spp. (milkweed) have been the subject of numerous ecological studies (Agrawal *et al.,* 2012) and digoxin, a cardenolide that is commercially isolated from *Digitalis* spp. (foxglove) (Kreis, 2017), is used for treating congestive heart failure and other diseases in both traditional and modern medicine.

Tissue-specific synthesis and transport of cardenolides have been studied most extensively in *Digitalis* spp. Although cardenolides occur in all *Digitalis* tissue types (Vogel and Luckner, 1981), not all tissues are capable of *de novo* cardenolide biosynthesis. Experiments with *Digitalis purpurea* cell cultures made from five different tissues showed that mesophyll cells have the greatest amount of cardenolide biosynthesis (Hagimori *et al.,* 1984). Root cells, which are not capable of *de novo* cardenolide biosynthesis (Hirotani and Furuya, 1977), nevertheless can modify cardenolides by hydroxylation, acetylation, and glycosylation (Reinhard, 1974). Experiments with *Digitalis lanata* showed that cardenolides with a terminal glucose in the oligosaccharide side chain, so-called “primary glycosides,” are actively taken up and stored in the vacuoles of *Digitalis* suspension culture cells, whereas those with other terminal sugars are not (Hoelz *et al.,* 1992; Kreis and Reinhard, 1987). Exogenous application of cardenolides to *D. lanata* leaves demonstrated that primary glycosides are transported both downward to the roots and upward to younger leaves (Christmann *et al.,* 1993). Uptake of cardenolides by *Macrosiphon rosae* (rose aphid) feeding on *D. lanata* indicated that the primary glycosides are transported in the phloem. By contrast, cardenolides were not found in the xylem sap (Christmann *et al.,* 1993).

The cardenolide-producing crucifer genus *Erysimum*, and in particular *Erysimum cheiranthoides* (wormseed wallflower), has been proposed as a genetic model system for studying cardenolide biosynthesis in plants (Munkert *et al.,* 2011, 2014; Züst *et al.,* 2018). Since the relatively recent (within the past 2-4 million years) evolution of cardenolide biosynthesis in *Erysimum,* there has been a rapid diversification of species in this genus (Züst *et al.,* 2020). An analysis of 48 *Erysimum* species by mass spectrometry identified almost 100 different cardenolide structures. Cardenolides in *E. cheiranthoides* have digitoxigenin, cannogenin, cannogenol, or strophanthidin as the steroid core, with further structural variation provided by the addition of sugar moieties (Züst *et al.*, 2020). Some of the more abundant *E. cheiranthoides* cardenolides, which are relevant to the current study, are illustrated in Figure 1.

**Figure 1.**
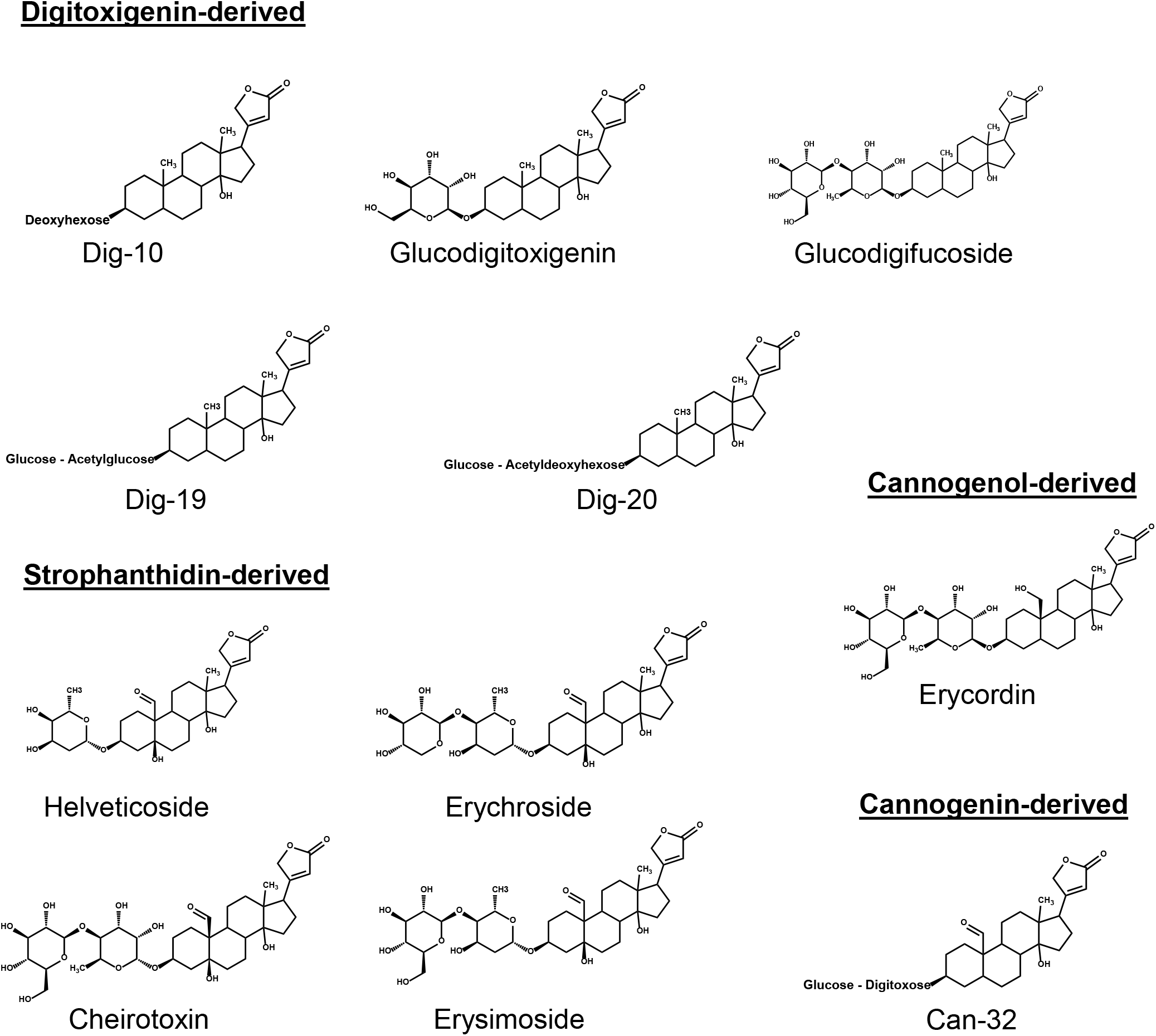
As described by Züst et al (2020), abundant cardenolides in *E. cheiranthoides* have digitoxigenin, cannogenol, cannogenin, or strophanthidin as the steroid core. Sugar side chains added to these steroid cores provide additional structural variation. The side chains of Dig-10, Dig-19, Dig-20, and Can-32 have not been fully characterized but are predicted based on MS fragmentation.

To investigate tissue-specific distribution and transport of cardenolides, we used *E. cheiranthoides* var. Elbtalaue, which has a sequenced genome and a well-characterized foliar cardenolide profile (Züst *et al.,* 2020), as well as mutant line 454 (Mirzaei *et al.,* 2020), an Elbtalaue derivative with a single-locus recessive mutation on chromosome 6 that causes elevated accumulation of digitoxigenin-derived cardenolides and lower abundance of cardenolides with cannogenol, cannogenin, and strophanthidin as the steroid core. The unambiguous differences in the cardenolide profiles of wildtype Elbtalaue and 454 mutant plants made it possible to investigate within-plant cardenolide transport by means of crosses and grafting experiments.

## 2. Results and Discussion

### 2.1 Tissue-specific variation in cardenolide abundance

To identify tissue-specific variation in cardenolide abundance, we analyzed *E. cheiranthoides* roots, whole above-ground plants, young leaves, mature leaves, old (senescing) leaves, buds, flowers, seed pods, dry seeds, and cotyledon-stage seedlings by HPLC-MS (Figure S1). Ten cardenolides with digitoxigenin, cannogenol, cannogenin, or strophanthidin as the steroid core were reliably identified in these assays. Principal component analysis (PCA; Figure 2) showed distinct cardenolide profiles in the different tissue types. Roots had a lower overall cardenolide content than aboveground plant parts (Figure S1). A similar observation has been made with *Asclepias syriaca* (common milkweed) (Rasmann *et al.,* 2009), and it was suggested that this is reflective of different herbivore pressure in aboveground and belowground plant parts. Consistent with the optimal defense theory (McKey, 1974; Meldau *et al.,* 2012), cardenolides, which had the highest concentrations in reproductive tissue and young leaves, became less abundant as leaves senesced (Figure S1). An exception was erychroside, which was almost completely absent from dry seeds (Figure S1B). Conversely, helveticoside was relatively abundant in dry seeds (Figure S1F). Notably, the cardenolide profile of seedlings at the cotyledon stage was most similar to that of dry seeds (Figure 2).

**Figure 2.**
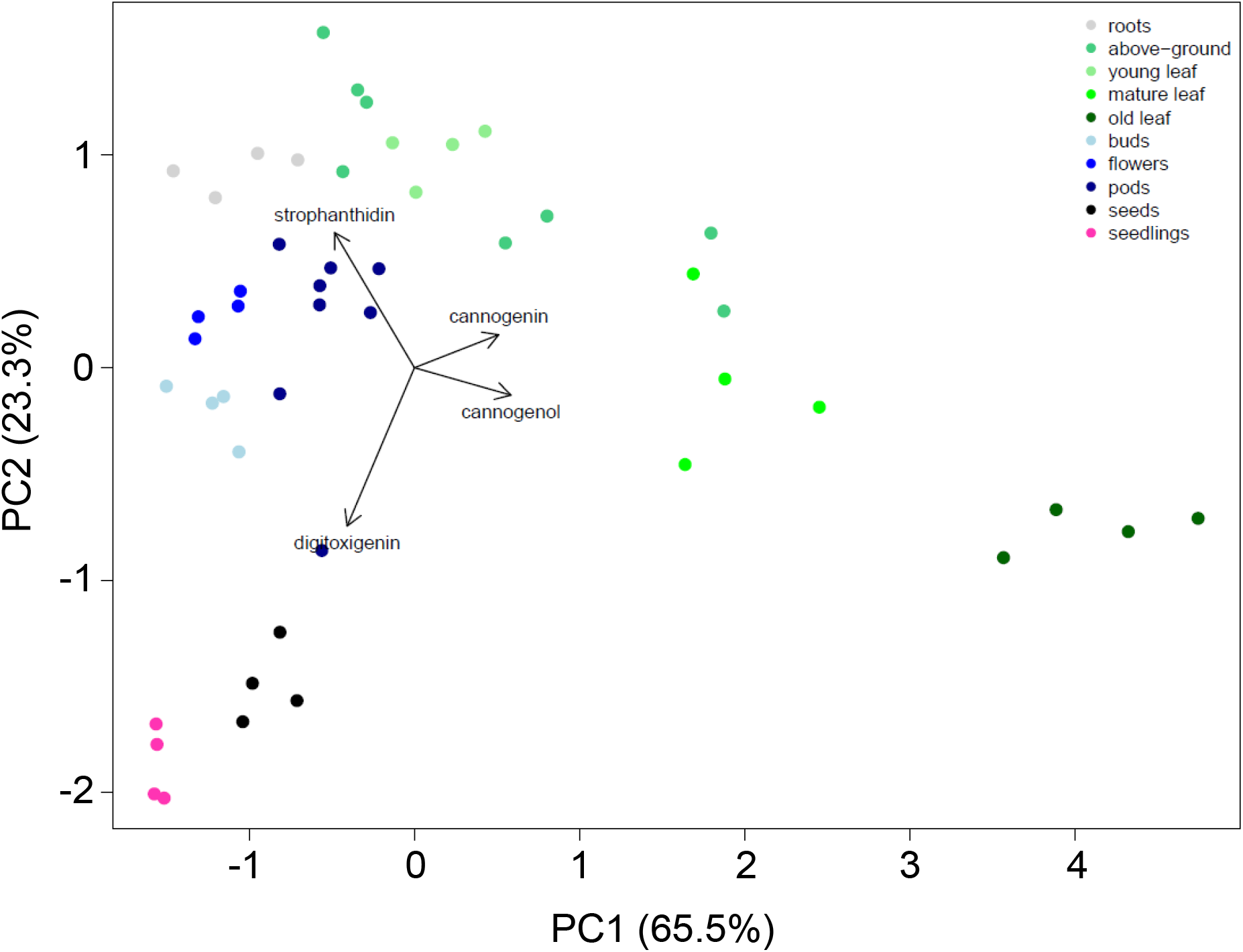
Principal component analysis (PCA) biplot of genin abundance as a percentage of total cardenolide abundance. Variable loadings for the first two principal components are displayed as vectors.

### 2.2 Cardenolide transport from maternal tissue to seeds

To determine whether there is transport to reproductive tissues, we measured the cardenolide content of mature dry seeds from crosses between wildtype *E. cheiranthoides* var. Elbtalaue and the 454 mutant line (Mirzaei *et al.,* 2020). These assays were done at a different time with a different column than the *E. cheiranthoides* tissue profiling in Figures 2 and S1, resulting in a slightly different but overlapping set of reliably detected cardenolides. The cardenolide profiles were similar in control experiments comparing seeds from naturally selfed plants and manual self-pollination (Figure 3), indicating that the manual pollination process did not have a significant effect on cardenolide accumulation. In reciprocal crosses between wildtype and 454 mutant plants, the seed cardenolide phenotype was determined entirely by the maternal genotype. Notably, in crosses where the maternal plant is 454 mutant and the pollen is wildtype (purple bars in Figure 3), the seeds nevertheless have the 454 cardenolide profile. Since the cardenolide phenotype of 454 mutation is recessive (Mirzaei *et al.,* 2020), this indicates that cardenolides are not synthesized in the seeds themselves, but rather originate from the maternal tissue.

**Figure 3.**
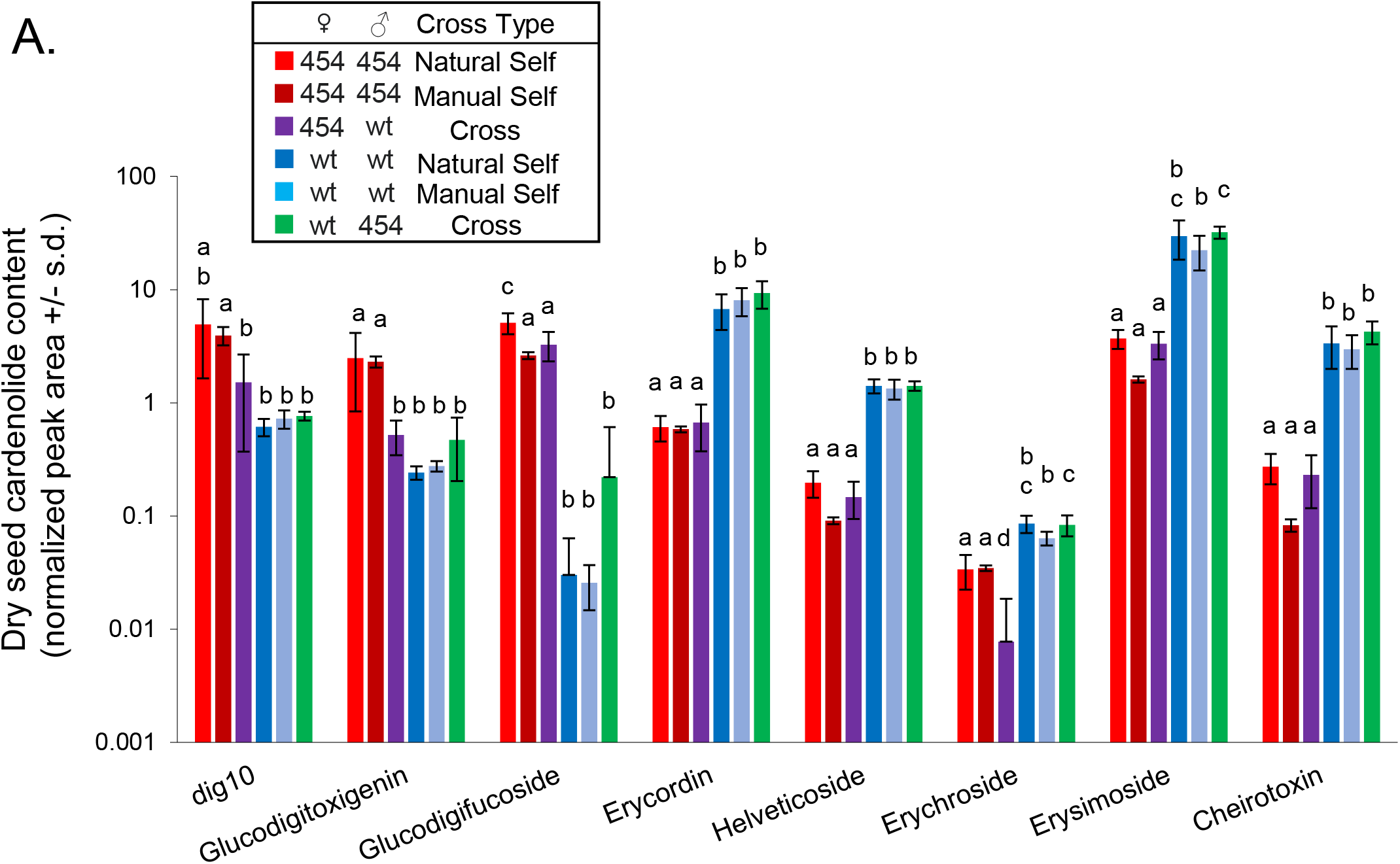
Maternal genotype determines seed cardenolide phenotype. Seed cardenolide content was measured in seeds from naturally self-pollinated and manually crossed plants. Different letters indicate P < 0.05 differences for each cardenolide, ANOVA followed by Tukey’s HSD test. Bars are mean +/− s.d. of N = 4-5. wt = wildtype *E. cheiranthoides* var. Elbtalaue, 454 = 454 cardenolide mutant line. Peak areas were normalized to a ouabain internal standard.

Given the similar cardenolide profiles of dry seeds and seedlings at the cotyledon growth stage (Figures 2 and S1), we hypothesized that the cotyledons contain cardenolides that were previously stored in the seeds. To test this hypothesis, we measured the cardenolide content of F1 seedlings from reciprocal crosses between 454 mutant and wildtype plants. Before true leaf emergence, the cardenolide content of the cotyledons was determined by the maternal plant genotype (Figures 4A and S2A). This may be indicative of maternal provisioning of progeny seeds, which are likely to germinate nearby and, at least initially, experience the same herbivore environment as their parent plant. However, once true leaves had emerged, the cardenolide phenotype of both cotyledons (Figure 4B and S2B) and true leaves (Figure 4C and S2C) was determined by the seedling genotype, with the 454 mutation being recessive to the wildtype allele. This indicates that *de novo* synthesis of cardenolides in seedlings is not initiated until the emergence of the first true leaves.

**Figure 4:**
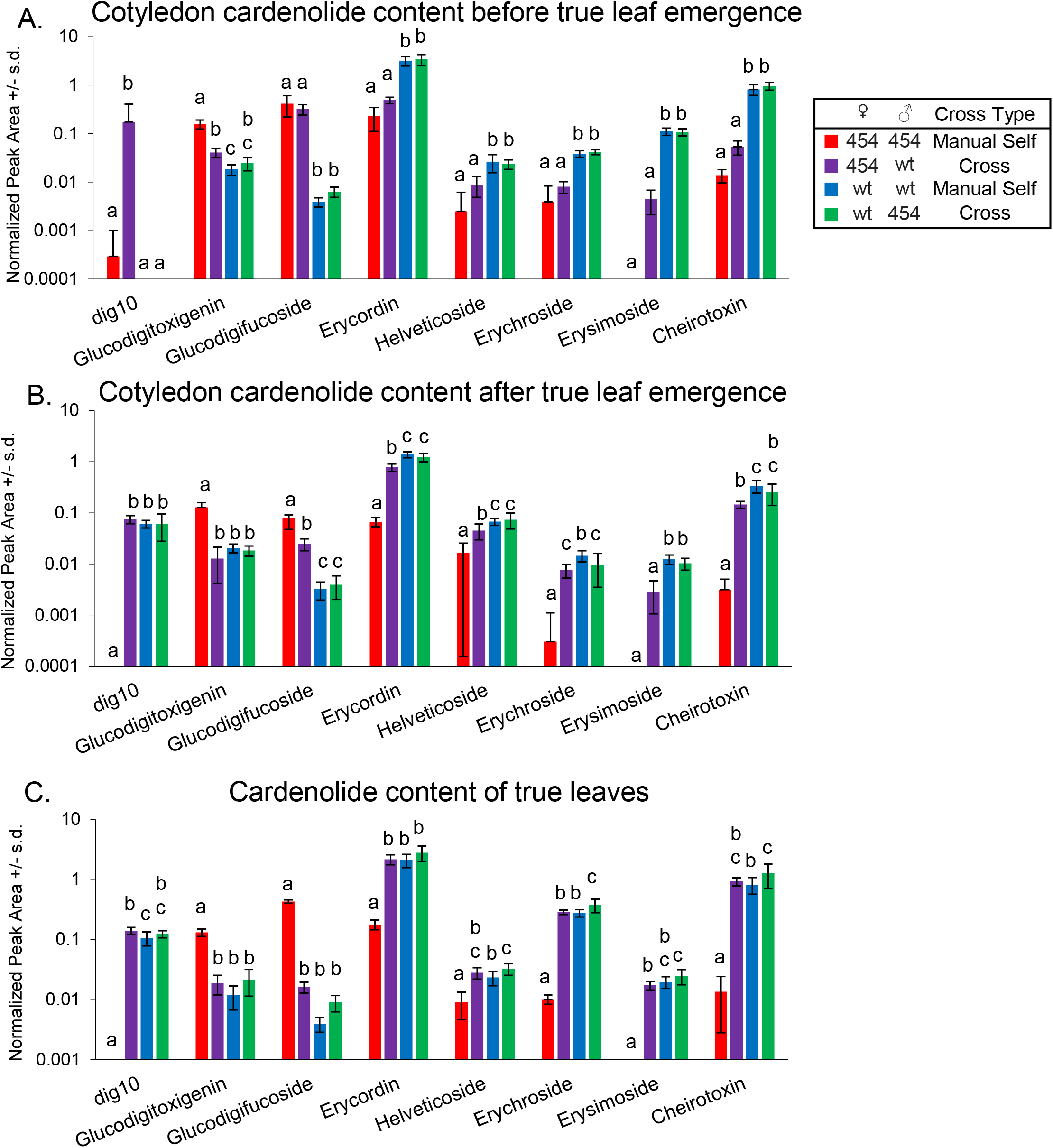
Cotyledons retain maternal plant cardenolide phenotype until true leaf emergence. (A) Cardenolide content of cotyledons before true leaf emergence, (B) cardenolide content of cotyledons after true leaf emergence, (C) cardenolide content of true leaves. Different letters indicate P < 0.05 differences for each cardenolide, ANOVA followed by Tukey’s HSD test. Bars are mean +/− s.d. of N = 6-10. wt = wildtype *E. cheiranthoides* var. Elbtalaue, 454 = 454 cardenolide mutant line. Peak areas were normalized to a ouabain internal standard.

Since the wildtype allele is dominant, we were not able to unequivocally conclude that cardenolide phenotype of F1 seeds from wildtype plants pollinated with 454 mutant pollen (green bars in Figure 3) was determined by the maternal genotype. To address this question, we measured the cardenolide content of single F2 seeds (Figure 5A) and seedlings (Figure 5B) from 454 x wildtype F1 plants. The genotype of these F2 seeds and seedlings was predicted to be segregating 1:2:1 homozygous wildtype : heterozygous : homozygous 454 mutant. However, when we assayed these seeds, they all had a wildtype cardenolide profile, indicating that the seed cardenolide is determined the by the maternal genotype, with homozygous 454 mutant seeds also having a cardenolide phenotype that is consistent with the maternal genotype. The segregating cardenolide genotype in the F2 generation was confirmed by letting the seedlings grow to the two-leaf stage. At this point, PCA showed two distinct cardenolide phenotypes in F2 seedlings, one that was similar to the wildtype and one that was similar to the homozygous 454 mutant (Figure 5C). Analysis of individual cardenolide abundances in F2 plants at a later growth stage (Figure 5D) showed both wildtype and 454 mutant profiles, indicating that at least some of the F2 seeds and cotyledons that were analyzed (Figure 5A,B) were homozygous mutant despite having a wildtype cardenolide phenotype.

**Figure 5:**
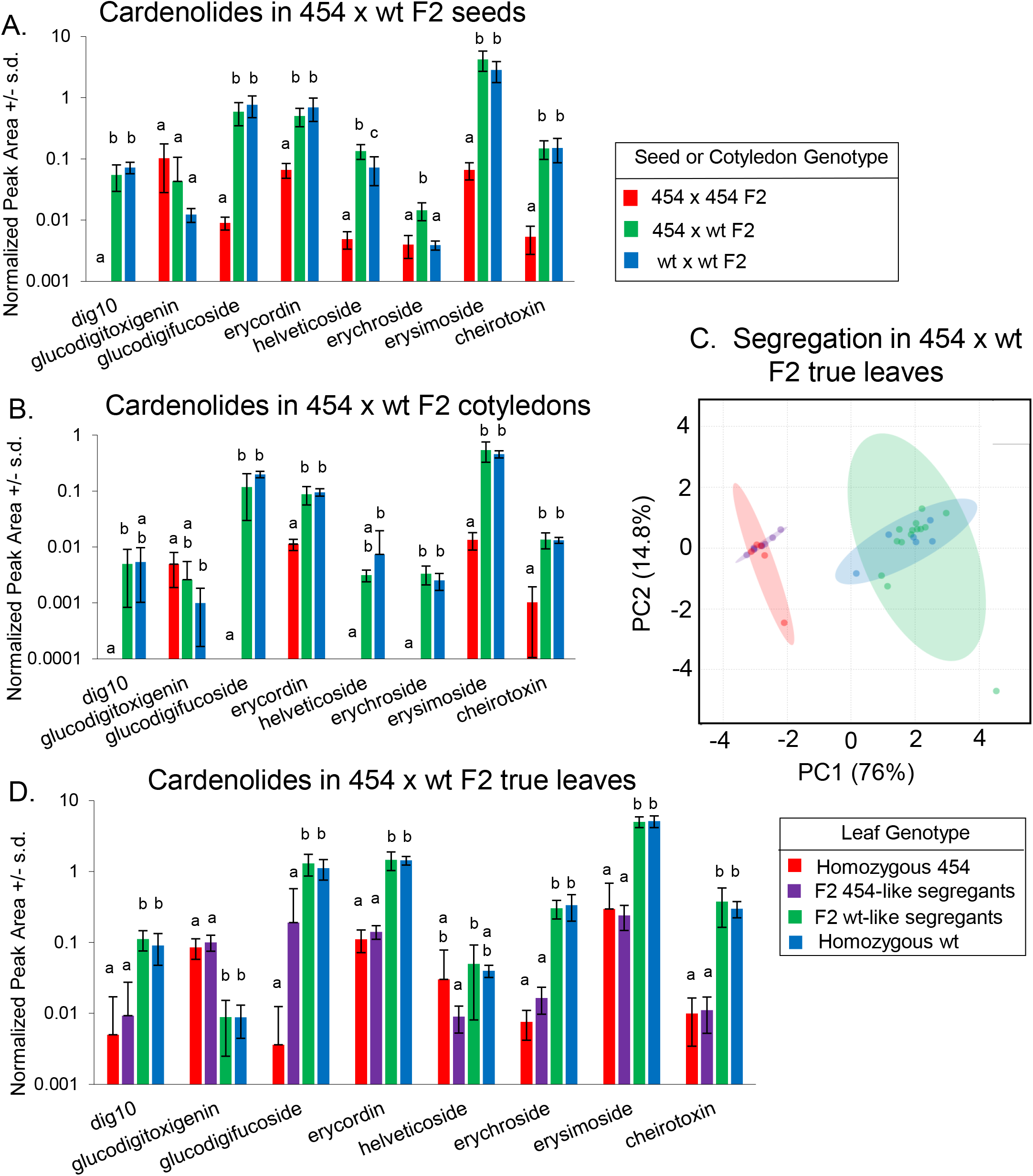
Cotyledons retain maternal plant cardenolide phenotype until true leaf emergence. Cardenolides were measured in F2 progeny of wt x 454 F1 plants. (A) Cardenolide content of single F2 seeds, (B) cardenolide content of F2 cotyledons from individual plants before true leaf emergence, (C) principal component analysis of cardenolide content of true leaves of F2 plants, and (D) cardenolide content of F2 true leaves. Different letters indicate P < 0.05 differences for each cardenolide, ANOVA followed by Tukey’s HSD test. Bars are mean +/− s.d. of N = 4-6 homozygotes and 20-24 F2s. wt = wildtype *E. cheiranthoides* var. Elbtalaue, 454 = 454 cardenolide mutant line. Ellipses in the PCA plot represent 95% confidence intervals. Peak areas were normalized to a ouabain internal standard.

### 2.3 Cardenolide transport from above-ground tissue to roots

The qualitative and quantitative differences in the cardenolide profiles of different *E. cheiranthoides* tissue types (Figures 2 and S1) could result from differences in localized biosynthesis or cardenolide transport from one plant organ to another. To investigate whether there is shoot to root cardenolide transport, we conducted reciprocal hypocotyl grafting experiments between wildtype and 454 mutant plants. Three weeks after moving successfully grafted plants to soil, we measured cardenolide content in the leaves and roots. The leaf cardenolide phenotype was consistent with the leaf genotype (Figure 6A). However, the root cardenolide phenotype was always determined by the leaf genotype, irrespective of whether the scion was wildtype or 454 mutant (Figure 6B). In *Digitalis* spp., cardenolides are transported to roots (Christmann *et al.,* 1993) but further modification can occur in root cells (Reinhard, 1974). By contrast, our results indicate that the root cardenolide profile in *E. cheiranthoides* is based on the transport of fully-formed cardenolides.

**Figure 6.**
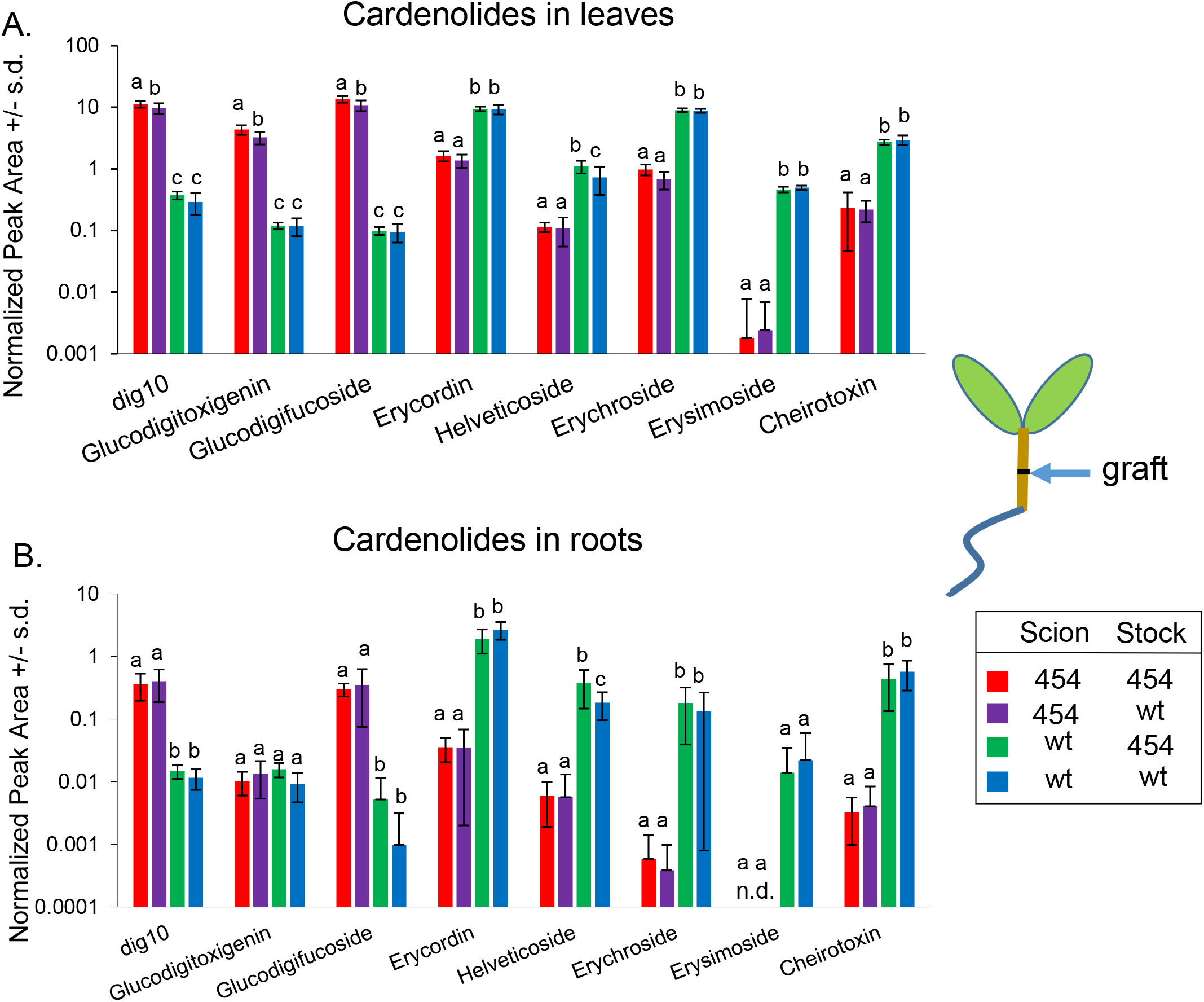
Grafting experiments show that shoot genotype determines root cardenolide phenotype. Seedlings were grafted at the cotyledon stage and cardenolides were measured in leaves and roots after three weeks. (A) Scion (leaf) cardenolides in grafted plants. (B) Stock (root) cardenolides in grafted plants. Different letters indicate P < 0.05 differences for each cardenolide, ANOVA followed by Tukey’s HSD test. Bars are mean +/− s.d. of N = 5-13 for shoot samples and 4-10 for root samples. wt = wildtype *E. cheiranthoides* var Elbtalaue; 454 = 454 cardenolide mutant line. Peak areas were normalized to a ouabain internal standard.

### 2.4 Cardenolide movement in vegetative tissue

To further delimit within-plant cardenolide movement, we conducted grafting experiments using three to four-week-old plants, such that the leaves above and below the graft junction have different genotypes. For control plants we made self-grafts where the stock and scion have the same genotype, either wildtype or 454 mutant. Two weeks after grafting, cardenolides were measured in leaves and stem segments from above and below the graft junction. PCA plots show that, both in leaves above the graft junction (Figure 7A) and in leaves below the graft junction (Figure 7B), the cardenolide phenotype is determined by the genotype of the measured leaf. This effect is consistent across all of the measured cardenolides (Supplemental Figure S3A,B), suggesting little or no leaf-to-leaf transport and that cardenolides are synthesized in leaf tissue. However, stem cardenolides in heterologously grafted plants had a more intermediate profile. This was the case both above the graft junction (Figures 7C and S3C) and below the graft junction (Figures 7D and S3D), indicating that cardenolides are transported both acropetally and basipetally. Further evidence for cardenolide transport in *E. cheiranthoides* phloem comes from the observation that these compounds are taken up by *Myzus persicae* (green peach aphids) (Mirzaei *et al.,* 2020).

**Figure 7.**
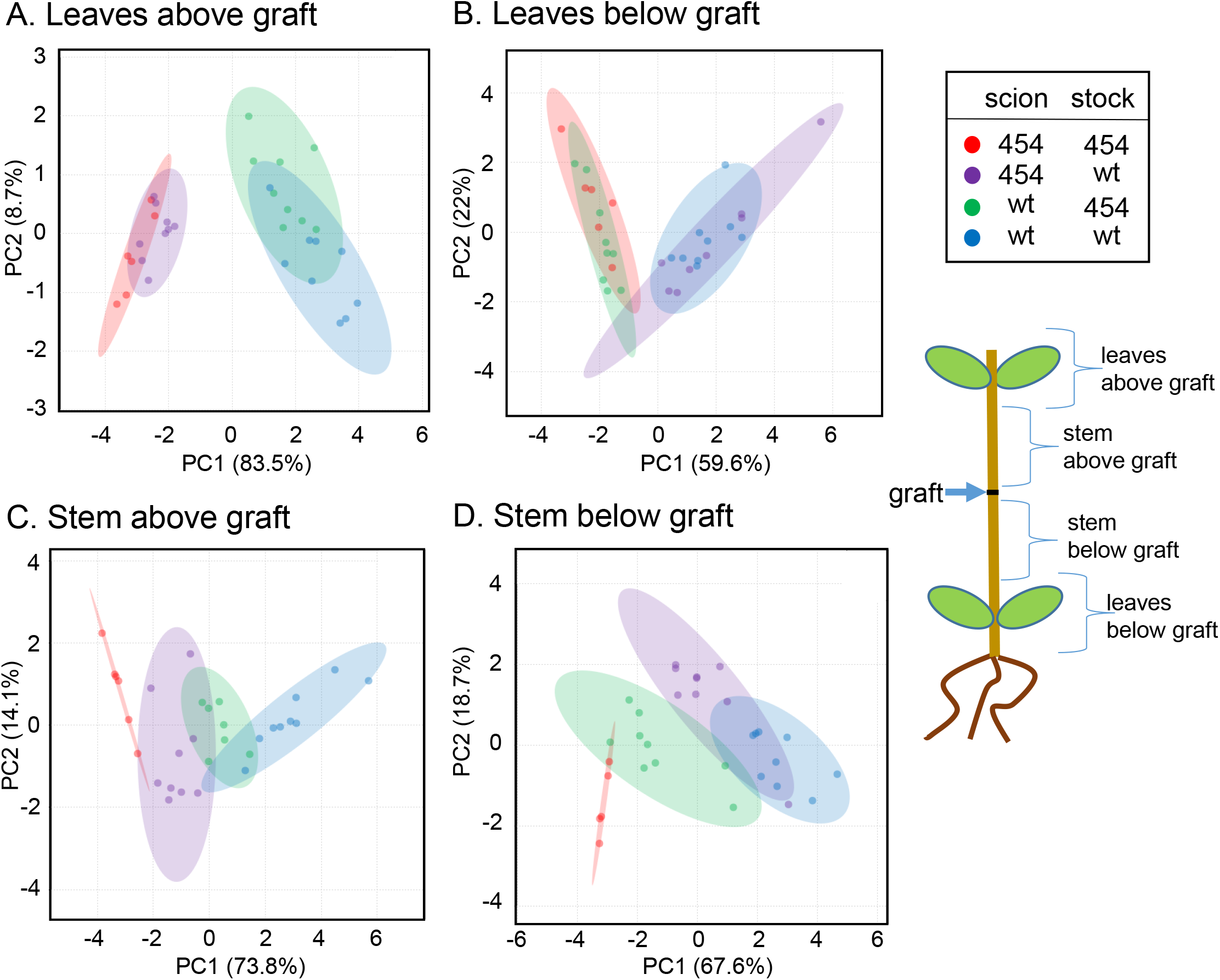
Cardenolides transport in aboveground tissue. The stalks of three to four-week-old wildtype and 454 cardenolide mutant plants were grafted. Two weeks later, cardenolides were measured in (A) leaves above the graft junction, (B) leaves below the graft junction, (C) stems immediately above the graft junction, and (D) stems immediately below the graft junction. The principal component analysis (PCA) is of eight detected cardenolides. Bar graphs of the data are in Supplemental Figure S3. Ellipses signify 95% confidence intervals.

Our crossing experiments (Figures 3) showed that seed cardenolide content is determined by the maternal genotype. However, this could be either due to “local” cardenolide transport *e.g.,* from the silique hulls to the seeds, or longer-distance transport from vegetative tissue. We used reciprocal grafts of wildtype and 454 mutant plants to determine whether the stock genotype affects the seed cardenolide phenotype. When inflorescences were grafted as scions on a stock with the same genotype, reproductive tissues had genotype-specific cardenolide profiles, as seen in the blue (wildtype) and red (454 mutant) samples in Figures 8 and S4. However, in the reciprocal inflorescence grafts of wildtype and 454 mutant plants, the flowers, green siliques, and dry seeds had intermediate cardenolide profiles, suggesting that there is not only transport of cardenolides from vegetative tissue to the inflorescences, but also more localized biosynthesis based on the inflorescence genotype.

**Figure 8:**
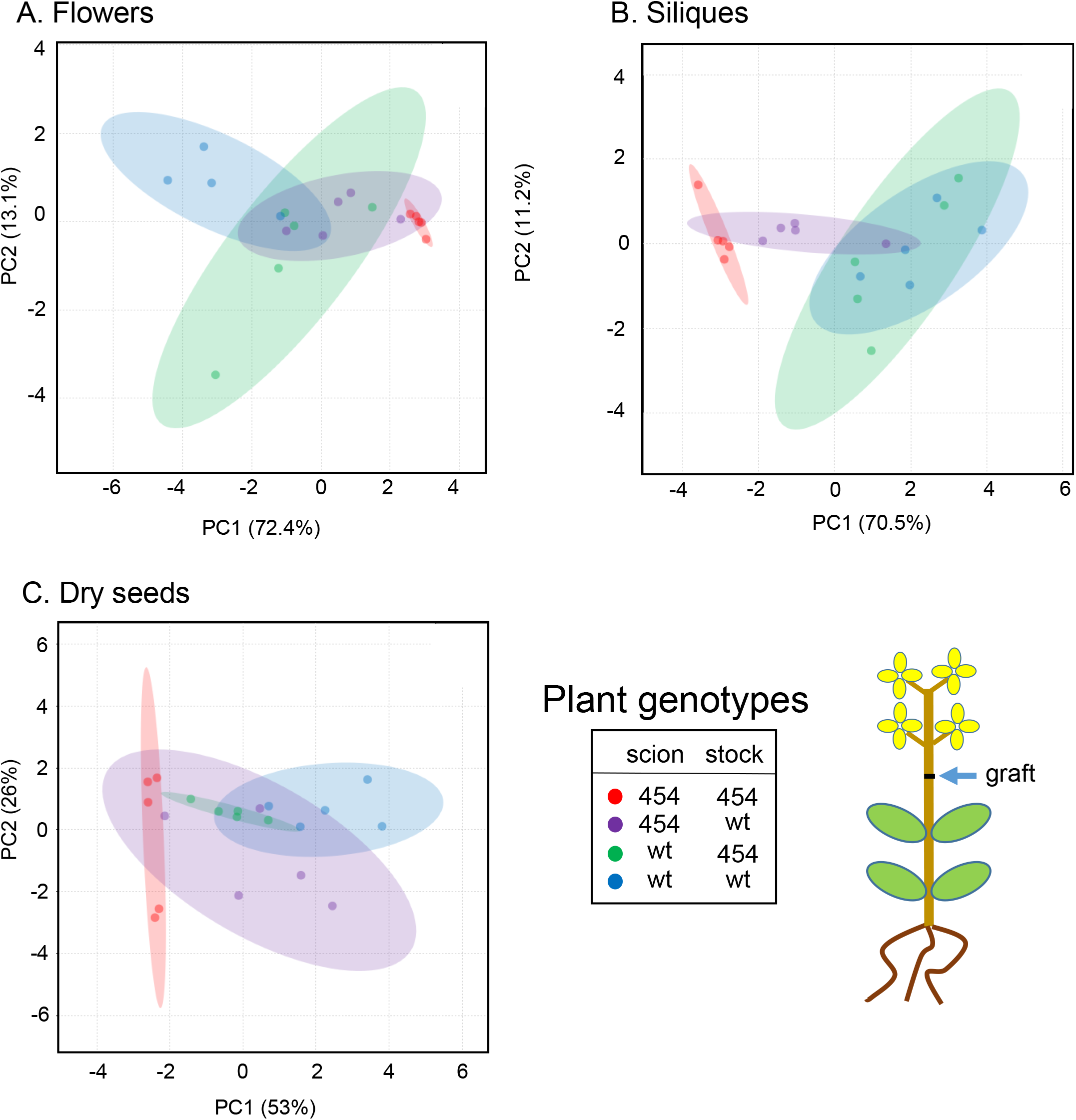
Intermediate cardenolide profiles in inflorescence-grafted plants. Plants were grafted at the inflorescence stage, with the graft junction above the leaves of the stock and below the developing inflorescence of the scion. Cardenolide content was measured in (A) flowers, (B) green siliques, and (C) dry seeds of the grafted plants. The principal component analysis (PCA) is of eight detected cardenolides. Ellipses signify 95% confidence intervals. Bar graphs of the data are in Supplemental Figure S4.

## 3. Conclusion

Our results provide new insight into the tissue-specific distribution and transport of cardenolides in *E. cheiranthoides.* In many cases, genes involved in the biosynthesis of plant specialized metabolites have been identified by correlating gene expression and metabolite abundance. However, extensive transport of cardenolides within *E. cheiranthoides* will complicate such research efforts. For instance, any correlative analysis to identify biosynthetic genes may have to assume that the expression levels in roots and developing seeds are very low or absent, despite the presence of cardenolides in these tissues. The transition from the cotyledon stage to the development of true leaves may be valuable for identifying differentially expressed genes that are upregulated at the time when new cardenolide biosynthesis is initiated.

Insects feeding on seeds, leaves, and roots of *E. cheiranthoides* are exposed to qualitative and quantitative variation in the cardenolide profiles. This is reflective of both the fitness value of these different tissue types and may also represent targeted defense against specific herbivore feeding guilds. Although we did not observe increased cardenolide accumulation in response to whole-plant defense induction with methyl jasmonate (Züst *et al.,* 2020), we cannot rule out reallocation of cardenolides to the sites of insect feeding. Future research will be required to identify the more specific defensive function of the diverse cardenolides that are present in *E. cheiranthoides*.

## 4. Experimental

### 4.1 Plant growth conditions

Experiments were performed using the genome-sequenced *Erysimum cheiranthoides* var Elbtalaue (Züst *et al.,* 2020), Arabidopsis Biological Resource Center (https://abrc.osu.edu) accession number CS29250, and the mutant line 454 (Mirzaei *et al.,* 2020). All soil-grown plants were grown in Cornell Mix (by weight 56% peat moss, 35% vermiculite, 4% lime, 4% Osmocote slow release fertilizer [Scotts, Marysville, OH], and 1% Unimix [Scotts]) in Conviron (Winnipeg, CA) growth chambers with 60% relative humidity, constant temperature of 23°C, and 180 μM m ^−2^ s^− 1^ photosynthetic photon flux density, and a 16:8 h day:night cycle. To promote germination, seeds were cold-stratified for two days at 4°C before planting.

Plants grown in axenic conditions for hypocotyl grafting were surface-sterilized as follows: 100-200 mg of *E. cheiranthoides* seeds were added to 1.5 ml microcentrifuge tubes and shaken in 70% ethanol for 1 min, the ethanol was removed by aspiration, a solution of 50% bleach and 0.05% Tween-20 in water was added, tubes were shaken for 1 min, the bleach solution was aspirated off, and seeds were washed four times in sterile deionized water. After suspension in sterilized 0.1% agar, seeds were cold-treated for two days at 4°C before planting on Murashige and Skoog (MS) agar plates [0.6% agar, ½ strength MS medium (Murashige and Skoog, 1962), and 0.5% sucrose]. Plates with seeds were placed in a Conviron growth chamber at 23°C, and 180 μM m ^−2^ s^− 1^ photosynthetic photon flux density, and a 8:16 h day:night cycle.

### 4.2 Sample collection for measurement of tissue-specific cardenolide content

Samples were collected from wildtype *E. cheiranthoides* for analysis of tissue-specific variation in cardenolide content: roots, whole above-ground plants, young leaves, mature leaves, senescing leaves, buds, flowers, seed pods, dry seeds, and seedlings. Dry seeds were harvested from mature plants, separated by sieving, and stored at 4°C until analysis. Seedlings were collected at the cotyledon stage, three days after germination on agar. Six to seven-week-old plants growing on soil were used for collecting leaves, buds, flowers, immature seed pods, and roots. Young leaves were newly emerged and about 1 cm long, mature leaves were fully expanded, and senescing leaves were starting to show signs of yellowing. Fresh plant samples were stored at −80°C until extraction and analysis by HPLC-MS as described previously.

### 4.3 Reciprocal crossing

An Arabidopsis crossing protocol (Weigel and Glazebrook, 2006) was adapted for *E. cheiranthoides*. Crosses were performed reciprocally between 454 and wild-type *E. cheiranthoides* plants (Figure S5) by removing sepals, petals, and stamens of unopened buds on the maternal plant using Dumont #5 tweezers, leaving only the exposed pistil. Multiple flowers from the paternal parent were removed and used to pollinate the maternal flower. Manual self-pollinations, as well as naturally self-pollinated plants, were produced as controls for each experiment.

### 4.4 Grafting

Hypocotyl grafting was performed using a protocol that was previously described for *Arabidopsis thaliana* (Arabidopsis) (Marsch-Martínez *et al.,* 2013), with modifications (Figure S5). Sterilized seeds were planted in Petri dishes containing agar medium (0.6% agar, ½ strength MS medium and 0.5% sucrose). The Petri dishes were sealed with Micropore tape (3M, St. Paul, MN) and placed in a growth chamber at a 60-degree angle (Figure S6A). Five-day-old seedlings were transferred to a cutting plate with a thin layer of 1% agar, where cotyledons were removed, and seedlings were cut in the middle of the hypocotyl using a #11 surgical blade (Feather Safety Razor Co., Osaka, Japan) (Figure S6B). The cut stock and scion were transferred to a recovery plate with the same MS agar formulation as described above and arranged such that the cut ends were touching. Grafted plants were returned to the growth chamber and left to recover for two weeks, after which grafting success was evaluated (Figure S6C) and successfully grafted plants were transferred to soil, where they grew for three weeks until tissue was collected (Figure S6D).

For grafting to produce plants with differing apical and basal shoot genotypes, plants were grown for 3-4 weeks until they were about 8 cm tall with well-developed rosettes. Stems were cut about 2 cm above the soil to create scion and stock samples (Figure S7A). To prevent desiccation after grafting, all except the smallest leaves were removed from the scion using a razor blade. Any leaves immediately below the graft junction on the stock were removed as well. The end of the scion was cut into a wedge shape and a short vertical cut was made in the stem of the stock using a razor blade, such that the scion wedge could fit evenly inside the cut stem. The graft junction was sealed with Parafilm (Bemis, Neenah, WI) and the entire flat of plants was covered with a plastic dome cover to maintain humidity until the graft junction healed (Figures S7B, S7C). Axillary shoots were removed to encourage apical shoot growth (Figure S7D).

For flower stalk grafting, wildtype and 454 mutant *E. cheiranthoides* plants were grown for seven weeks until inflorescences developed. About 5 cm of the apical flower stalk, lacking leaves, was removed using a razor blade and all flowers, buds, and siliques were removed except for the very terminal buds (Figure S8B). Siliques immediately below the graft junction were removed from the stock plant (Figure S8A). Grafting was performed as described above for rosette-stage plants, and grafted flower stalks were covered with plastic bags to prevent desiccation until the graft junctions healed (Figures S8A and S8B). Axillary flower stalks were removed to encourage growth of the grafted shoot (Figure S8C). Grafted plants were allowed to recover for two weeks (Figure S8D) before being harvested for cardenolide assays of flowers and siliques. Non-grafted control plants were left to mature and dry seeds were collected for assays.

### 4.5 Seedling cotyledon collection

Reciprocal F1 seeds from crosses between 454 and wild-type plants, as well as manually self-pollinated seeds from both genotypes, were cold-treated for 2 d and planted in soil. Each pollination event was considered a biological replicate. Seeds from individual siliques were grouped together when planting and tissue from multiple plants from the same silique was grouped together into a single sample for analysis. Samples were collected from seedlings at two time points, before and after true leaf development. At the first time point, 4-5 day old seedlings were harvested and their cotyledons were isolated and collected. For the second time point, 7-8 day old seedlings were harvested and their cotyledons and true leaves were collected separately. All samples were flash-frozen in liquid nitrogen and stored at −80°C until extraction.

### 4.8 Plant Extracts for UHPLC-MS

Plant tissues from grafting and leaf infiltration experiments (30-60 mg) were collected, weighed, and flash frozen in liquid nitrogen in 1.5 ml microcentrifuge tubes. Three 3 mm stainless steel balls were added to each tube, and tissues were homogenized in a Harbil 5G-HD shaker (Fluid Management, Wheeling, IL) for two minutes. Three volumes of 70% methanol containing 15 mg/L of either ouabain or digitoxin as an internal standard was added to each sample and vortexed to mix. Homogenized plant tissue was centrifuged at 13,500 g for 10 minutes to remove particulates, and if necessary, was transferred to a new tube and centrifuged a second time to ensure removal of particles. The resulting supernatant was transferred to glass vials with inserts (Thermo Scientific, Waltham, MA) and stored at −20C until analysis.

Due to the small size of *E. cheiranthoides* seeds and seedlings, an alternative method of extraction was performed to obtain a sufficient volume of extract for UPLC analysis. One or five dry seeds were placed in 2 ml microcentrifuge tubes and homogenized using plastic pestles. After addition of 100 μl of 70% methanol, samples were vortexed to mix, and seed debris was pelleted by centrifugation at 13,500 g for 4 min. Cotyledons and true leaves from reciprocal crossing experiments were collected in 1.5 mL tubes and homogenized using stainless steel balls and a paint shaker, as described above. After addition of 100 μl 70% methanol, samples were vortexed to mix, and the mixture was centrifuged at 13,500 g for 10 min. For both sample types, the supernatant was transferred to new 1.5 ml tubes and dried down in a Savant Speed Vac SC110 vacuum concentrator (Thermo Scientific) at medium heat setting, then re-dissolved in a volume of 70% methanol containing 15 mg/L ouabain internal standard proportional to each sample’s mass. Extracts were transferred to glass vials and stored at −20°C until analysis.

### 4.9 UHPLC-MS Analysis

Samples were separated using a Dionex UltiMate 3000 UHPLC (Thermo Scientific) equipped with a Titan C18 UHPLC column (100 × 2.1 mm, 1.9μm; Supelco Analytical, Bellfonte, PA) and analyzed with a Q-Exactive hybrid quadrupole-orbitrap mass spectrometer (Thermo Scientific). Two μl injections of samples were subjected to the following linear solvent gradient: for 1.5 seconds, the solvent mixture was held at 98% mobile phase A (water +0.1% formic acid) and 2% mobile phase B (acetonitrile +0.1% formic acid). Mobile phase B increased from 2% to 97% over 9.5 minutes, and was held at 97% for 1.5 minutes, before returning to 2% for 1.5 minutes. The flow rate was maintained at a constant 0.5 ml/min, the column was heated to 40°C, and the samples were kept at 15°C. Eluted compounds were detected from 0.6 to 13 minutes with a mass range of 150 to 900 m/z and with 140,000 resolution (full width at half-maximum) in positive ionization mode.

### 4.10 MS Data Analysis

Cardenolides were identified from positive-ion chromatograms and confirmed with fragmentation patterns common to cardenolides. Masses and retention times of the identified cardenolides a listed in Supplemental Table S1. Some cardenolides had very low abundance in tissues and were more easily detected by their sodium adducts. The sodium adducts of cheirotoxin, erysimoside, and erycordin were used for quantification in the crosses and grafting experiments. For measurements of tissue-specific cardenolide content, all fragment peaks were summed for quantification.

Raw file outputs were converted to mzML format using MSConvertGUI 64-bit (ProteoWizard 3.0.18239 64-bit) with the following settings: Output format: mzML; Binary encoding precision: 64-bit; Filters: Peak Picking. The output mzML files were entered into a peak detection pipeline based in R 4.0.1 (R Core Team 2020). Peak detection was performed using XCMS (Benton *et al.,* 2010; Smith *et al.,* 2006; Tautenhahn *et al.,* 2012, 2008) with the following parameters: the centWave method was used with an allowed error rate of 5 ppm, an allowed width range for chromatogram peaks of 3-15 seconds, a signal to noise ratio cutoff of 4, and an m/z cutoff value for dividing peaks with the same retention time of 0.01. Raw data were used for integration, and pre-filter requirements specified that a mass trace must have at least 3 peaks with an intensity value of at least 100 to be included. Peak grouping was performed as follows: the peak density method was used with a bandwidth value of 2, a width of overlapping m/z slices for peak grouping of 0.025, a minimum required sample fraction of 0.1, a minimum number of samples of 1, and a maximum number of groups of 50. Peaks were grouped using CAMERA (Kuhl *et al.,* 2012). Initial groups were made using 20% of the full width at half maximum. Isotopes were grouped with an allowed error of 5 ppm, an absolute error cutoff of 0.002 Daltons. Only one isotope peak was allowed, the minimum fraction of samples required to have the correct C12/C13 isotope ratio was 0.9, the correlation threshold for peak grouping was 0.7, and the p-value threshold for testing correlation significance was 0.1. In the resulting csv file, individual cardenolide compounds were manually identified using masses and known retention times listed in Supplemental Table S1.

### 4.11 Statistical analysis

Principal component analyses in Figures 5, 7, 8, and S2 were performed in Metaboanalyst 5.0 (Pang *et al.,* 2021) using default settings and auto scaling. All other statistical comparisons were conducted in R 4.0.1 (R Core Team, 2020) using functions available in the base stats package. The PCA for tissue-specific cardenolide abundances in Figure 2 was created using the function prcomp, with variables scaled to have unit variance. Compounds were normalized to the total cardenolide abundance in each sample and compound abundances were aggregated by genin. Raw data underlying all graphs are included in Supplemental Table S2.

## Acknowledgements

This research was funded by US National Science Foundation award 1645256, United States Department of Agriculture – National Institute of Food and Agriculture award 2020-67013-30896, and a Triad Foundation grant to GJ; a Summer Undergraduate Research Fellowship from the American Society of Plant Biologists and a Rawlings Cornell Presidential Research Scholar award to MLA; and a Cornell Chemistry Biology Interface Training Program (National Institute of Health/National Institute of General Medical Sciences award 5T32GM008500) fellowship and a US National Science Foundation Graduate Research Fellowship under Grant No. DGE-1650441 to GCY.

---

**Supplemental Figure S1.**
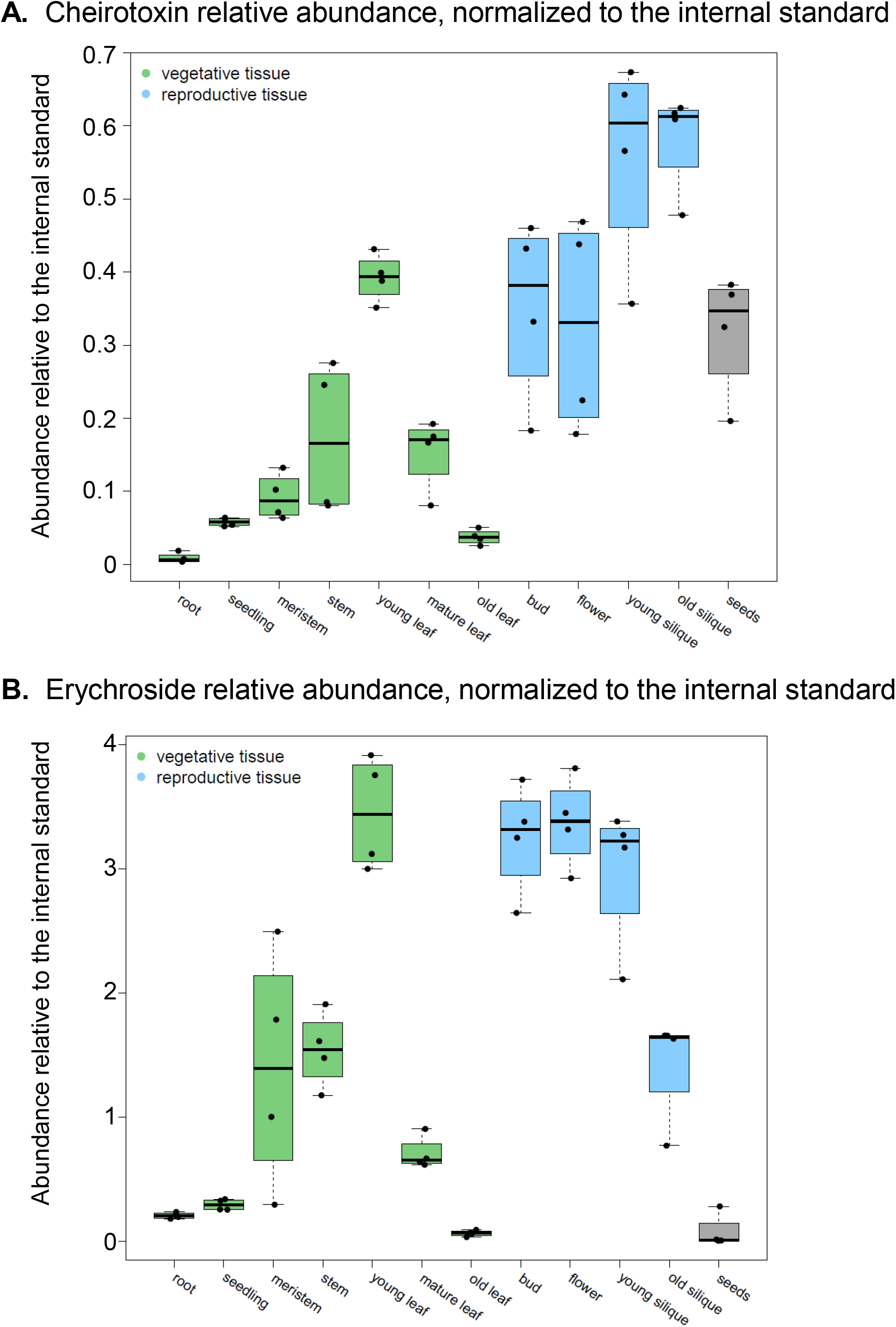
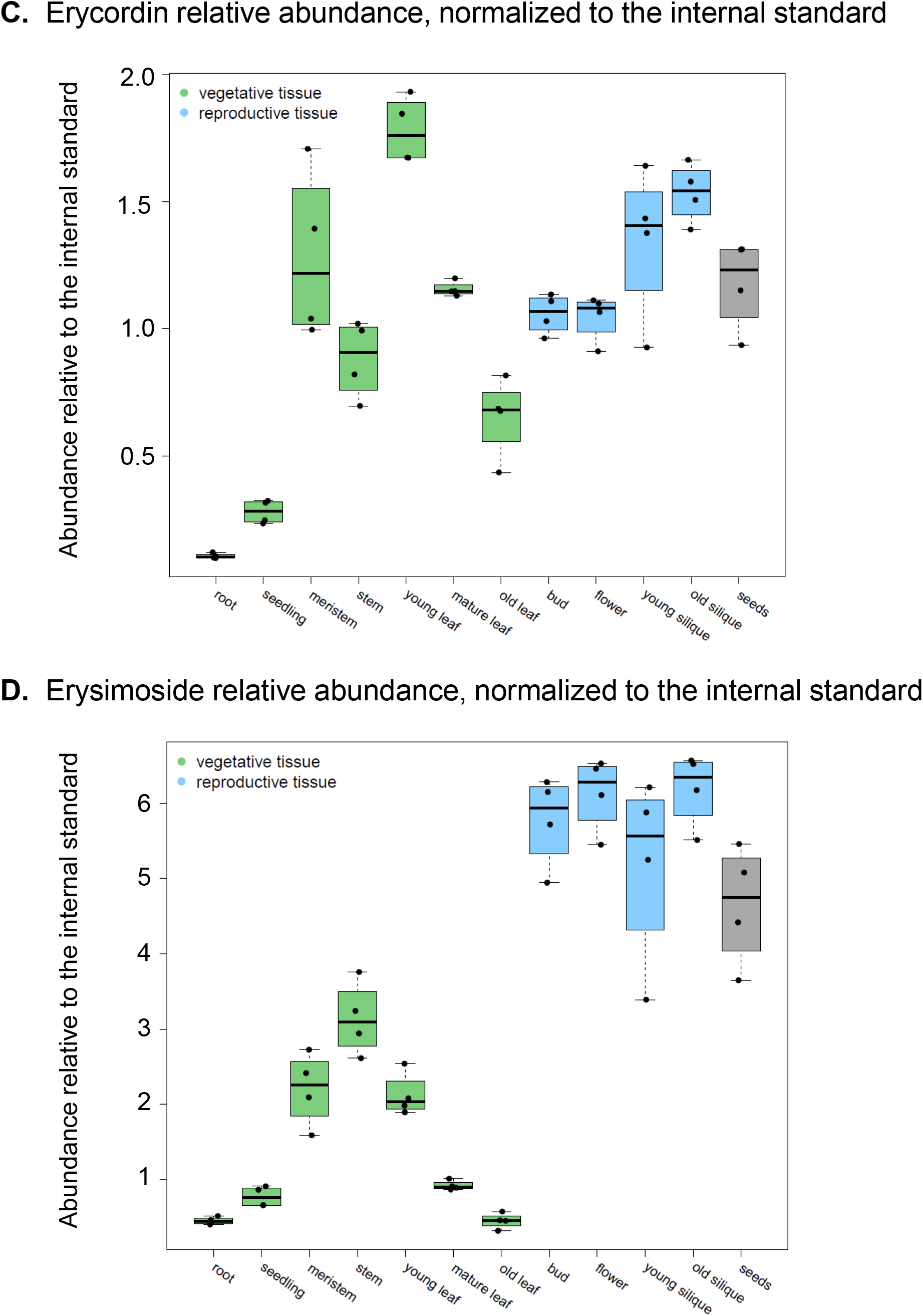
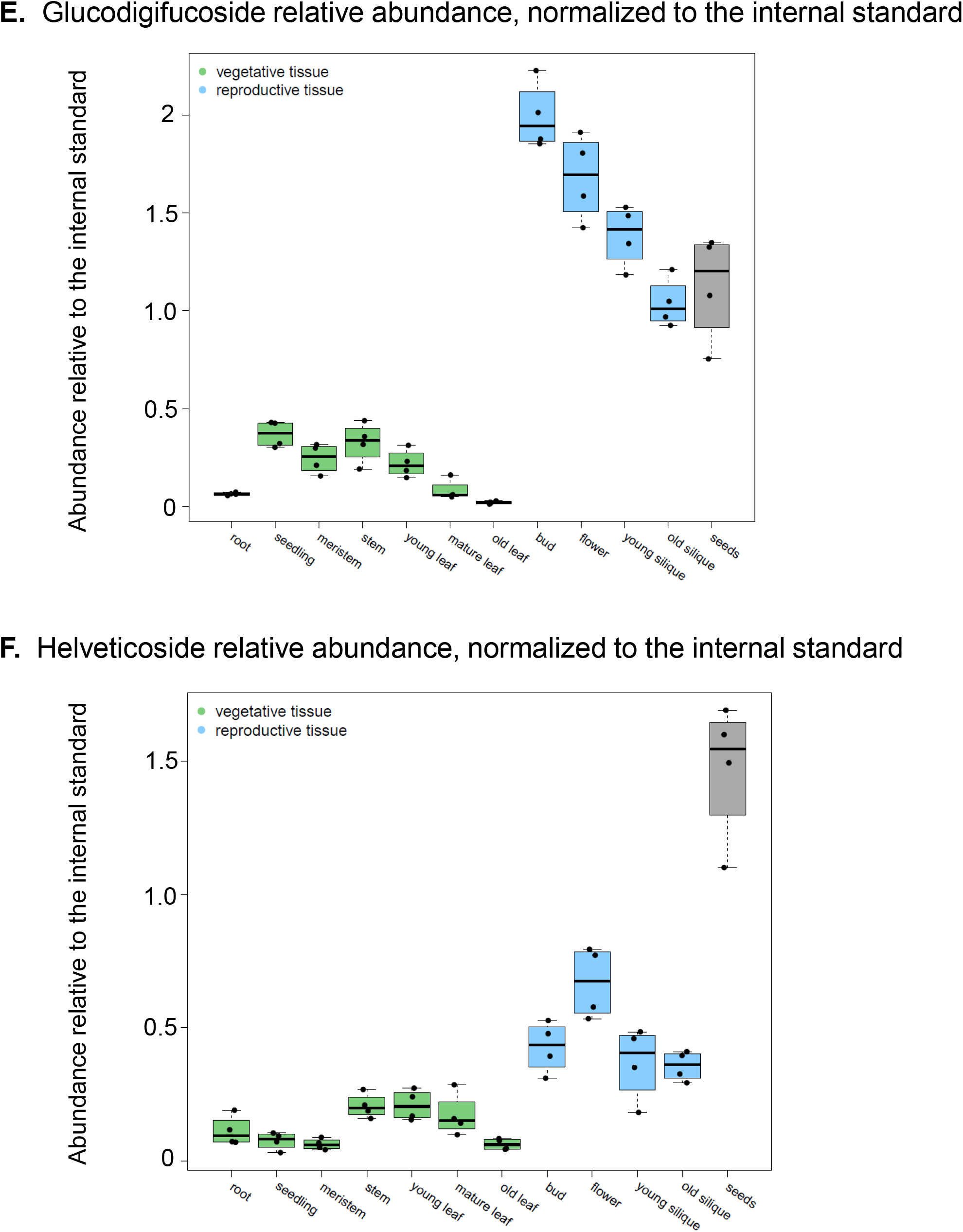
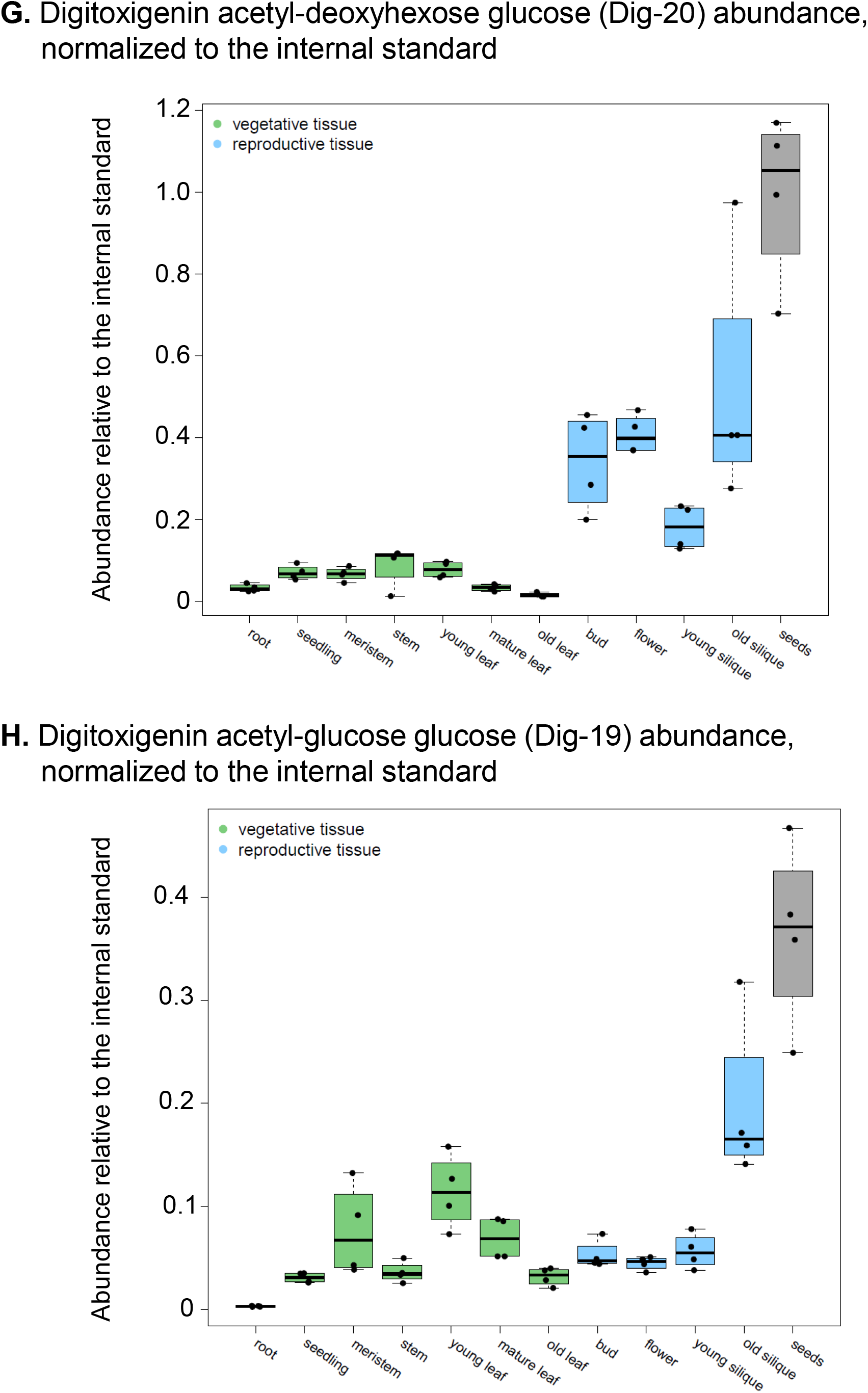
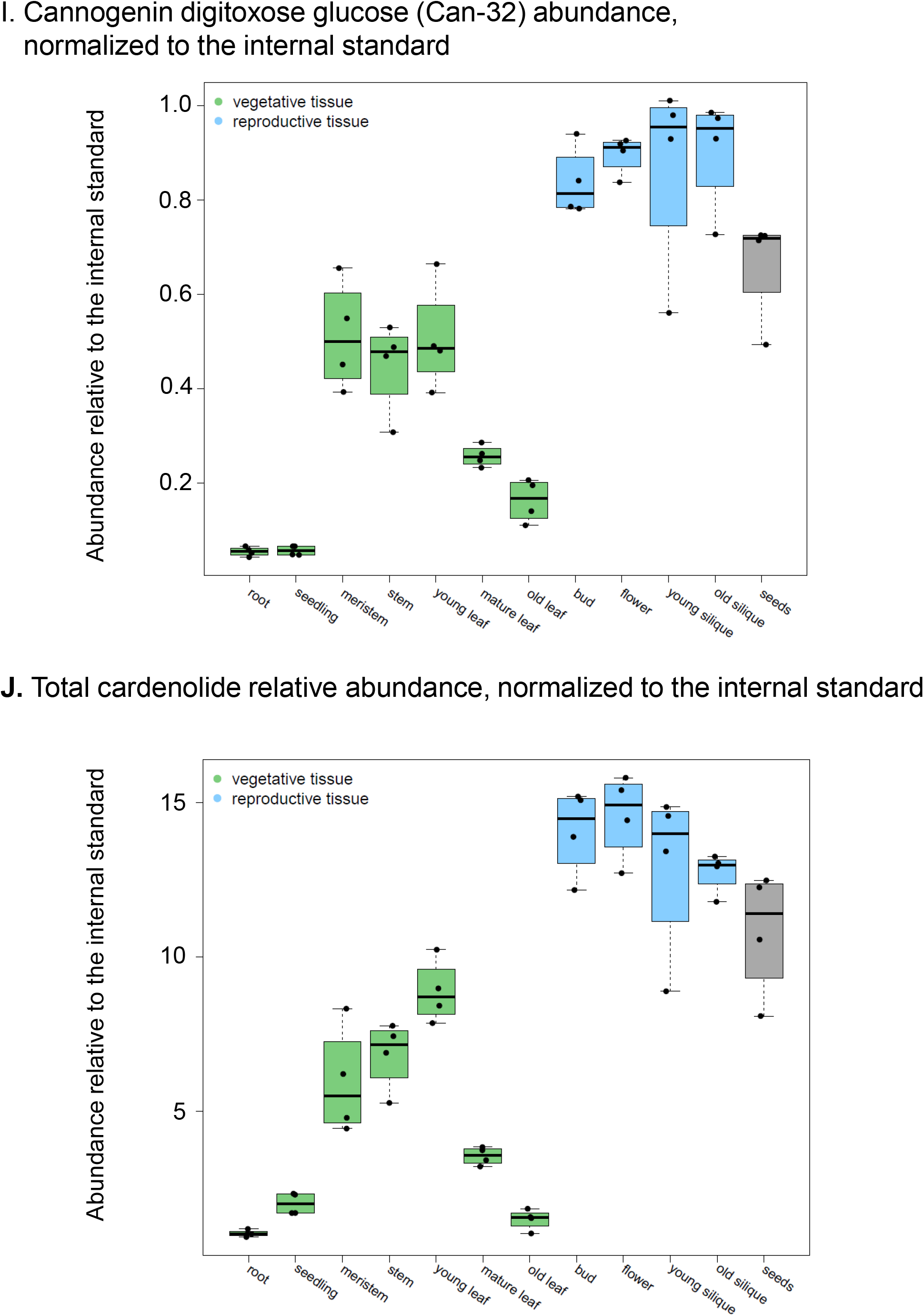
Relative cardenolide content in different *E. cheiranthoides* tissue types. (A) cheirotoxin (B) erychroside, (C) erycordin (D) erysimoside (E) glucodigifucoside, (F) helveticoside, (G) Digitoxigenin acetyl-deoxyhexose glucose (Dig-20), (H) Digitoxigenin acetyl-glucose glucose (Dig-19), (I) Cannogenin digitoxose glucose (Can-32), and (J) total cardenolide peak area relative to the digitoxin internal standard. Cardenolides in roots, whole above-ground plants, young leaves, mature leaves, old (senescing) leaves, buds, flowers, seed pods, dry seeds, and cotyledon-stage seedlings were measured by HPLC-MS. Four samples of each tissue type were measured. The data are represented as box plots showing the median, interquartile range, maximum and minimum after removal of outliers, and individual data points.

**Supplemental Figure S2:**
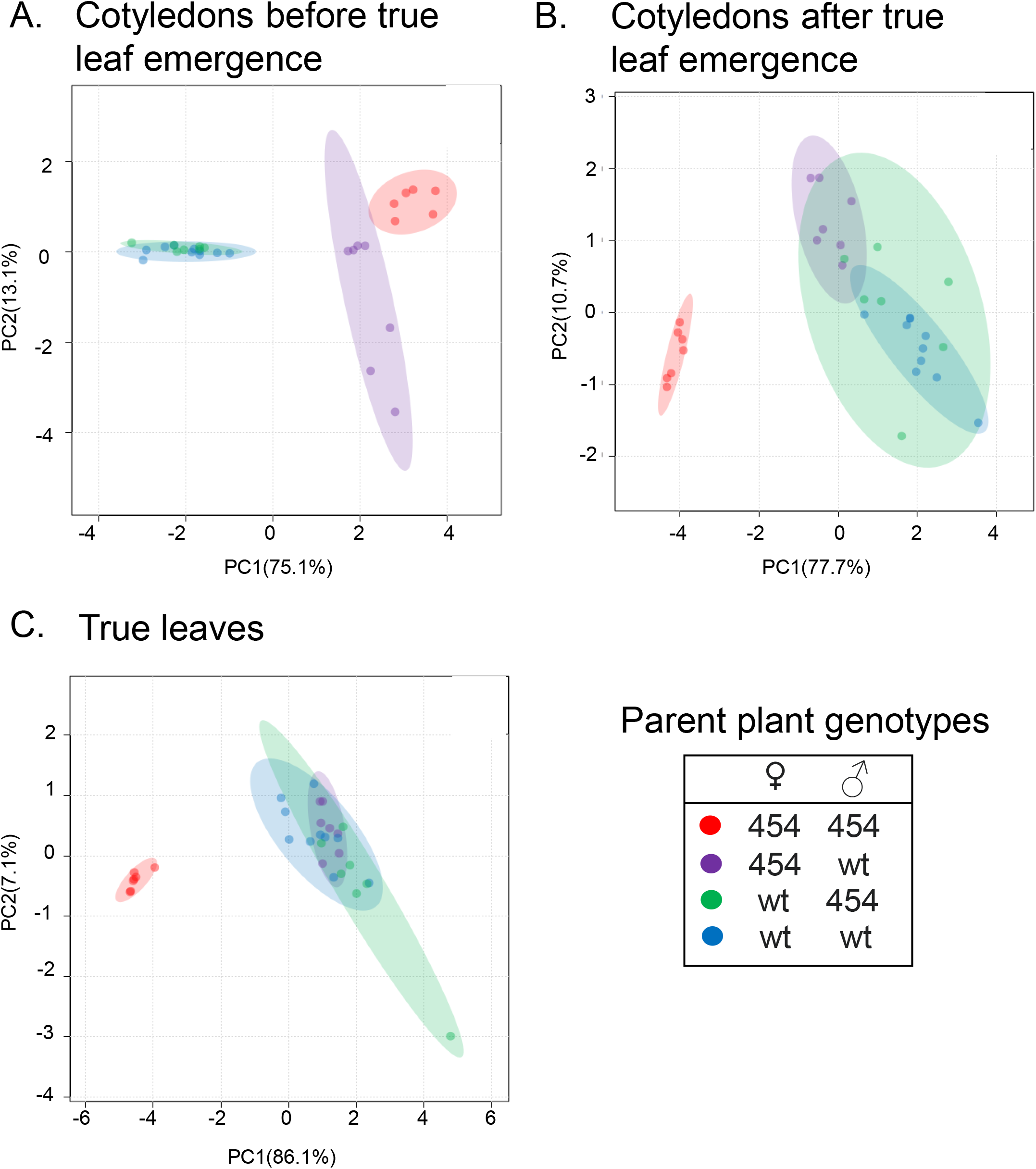
Cotyledons retain maternal plant cardenolide phenotype until true leaf emergence. The plots show principal component analysis (PCA) of the cardenolide data shown in Figure 4. (A) Cardenolide content of cotyledons before true leaf emergence, (B) cardenolide content of cotyledons after true leaf emergence, (C) cardenolide content of the first true leaves. The ellipses in the graphs represent 95% confidence intervals.

**Supplemental Figure S3:**
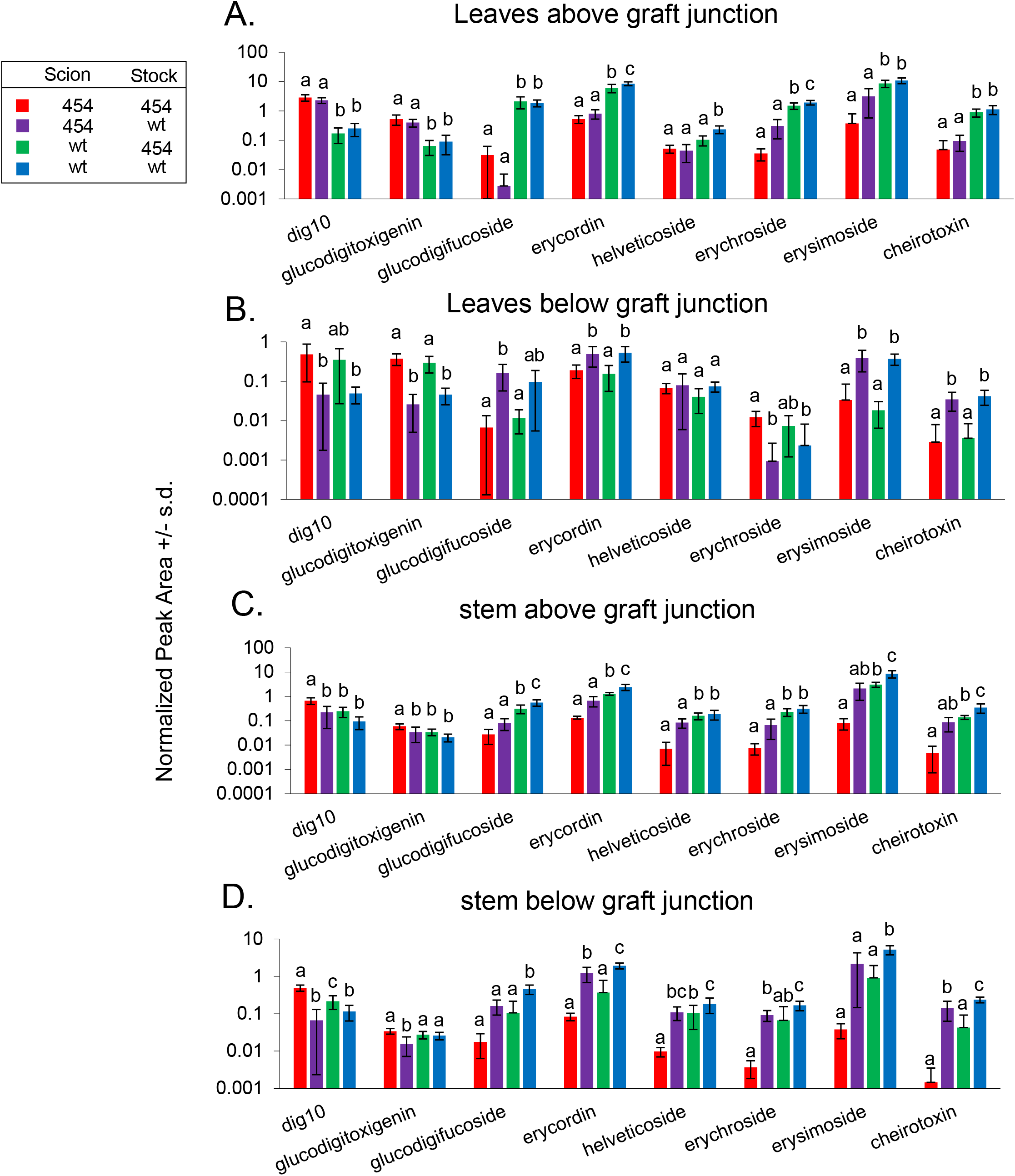
Leaves retain native cardenolide profiles in juvenile-grafted plants. Cardenolides were measured in five-week-old grafted plants from 4 different tissues: (A) upper leaves, (B) lower leaves, and stem tissues directly above and below the graft junction (C) and (D) respectively. Different letters indicate P < 0.05 differences for each cardenolide, ANOVA followed by Tukey’s HSD test. Bars indicate mean +/− s.d. of n= 5-9 plants. wt = wildtype *E. cheiranthoides* var. Elbtalaue, 454 = 454 cardenolide mutant line. Peak areas were normalized to a ouabain internal standard.

**Supplemental Figure S4:**
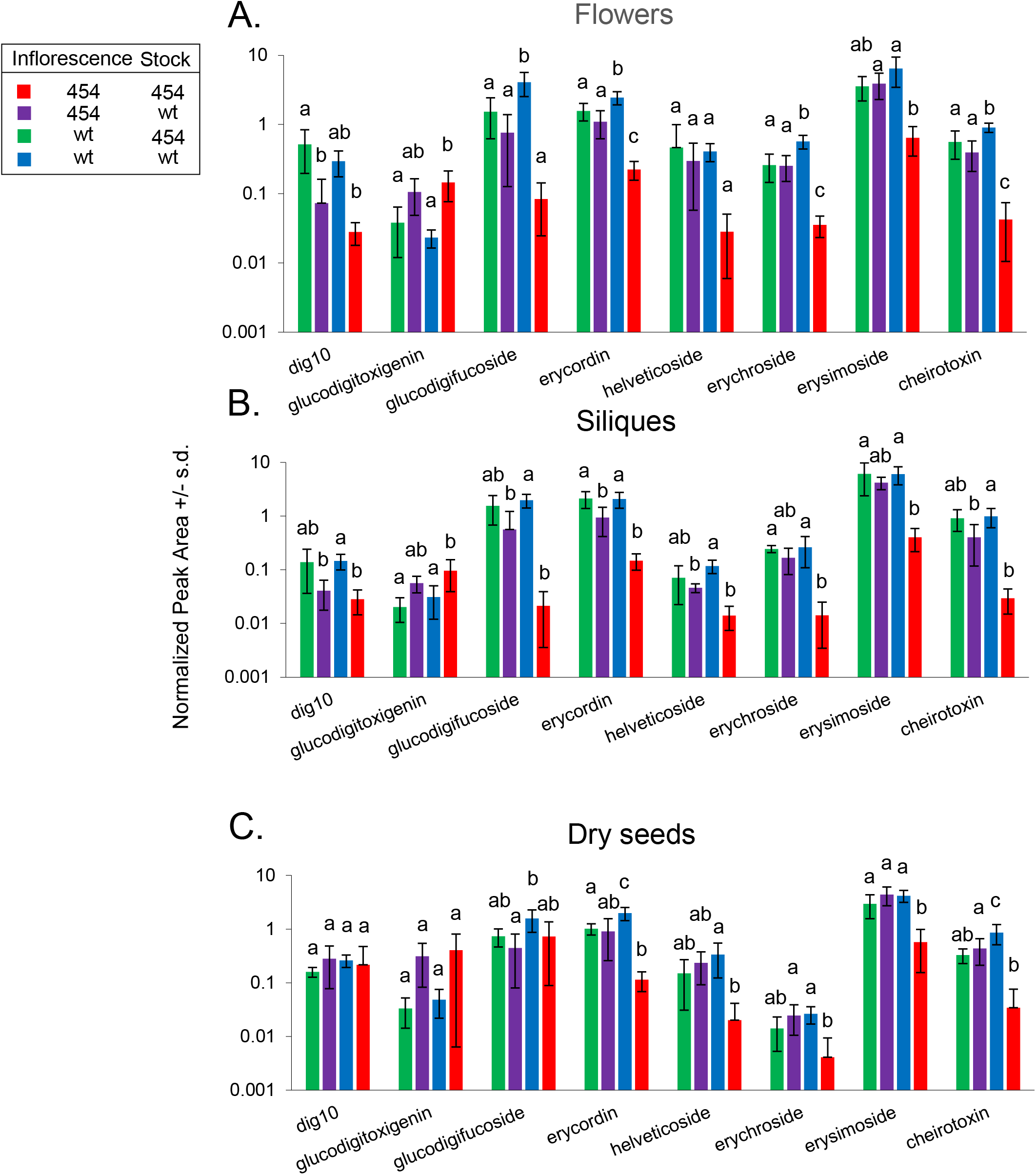
Intermediate cardenolide profiles in inflorescence-grafted plants. Plants were grafted at the inflorescence stage, with the graft junction above the leaves of the stock and below the developing inflorescence of the scion. Cardenolide content was measured in (A) flowers, (B) green siliques, and (C) dry seeds of the grafted plants. Different letters indicate P < 0.05 differences for each cardenolide, ANOVA followed by Tukey’s HSD test. Bars indicate mean +/− s.d. of n= 4-5 plants. wt = wildtype *E. cheiranthoides* var. Elbtalaue, 454 = 454 cardenolide mutant line. Peak areas were normalized to a ouabain internal standard.

**Supplemental Figure S5.**
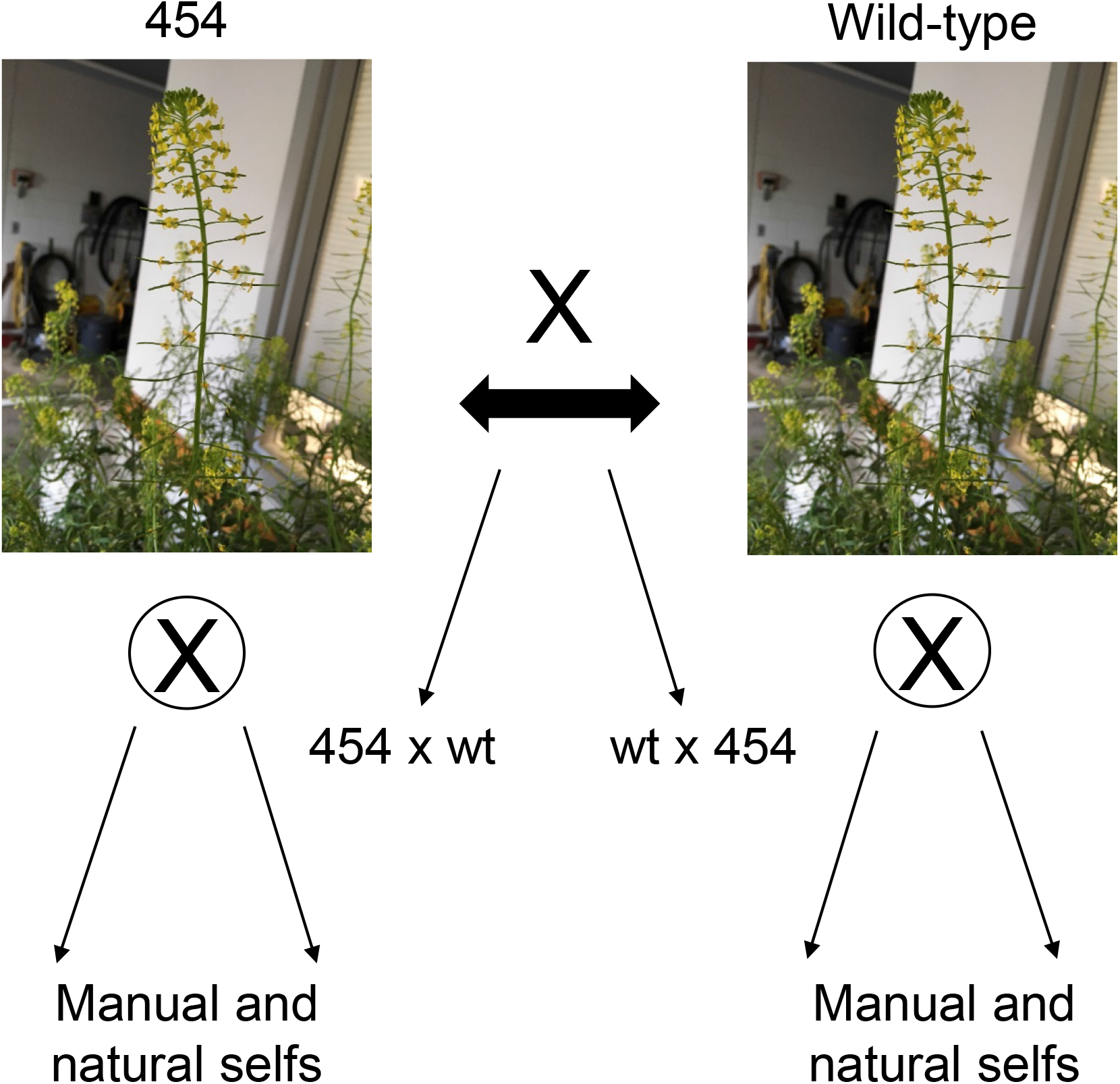
Crossing scheme to generate seeds with a different genotype than the maternal plant.

**Supplemental Figure S6.**
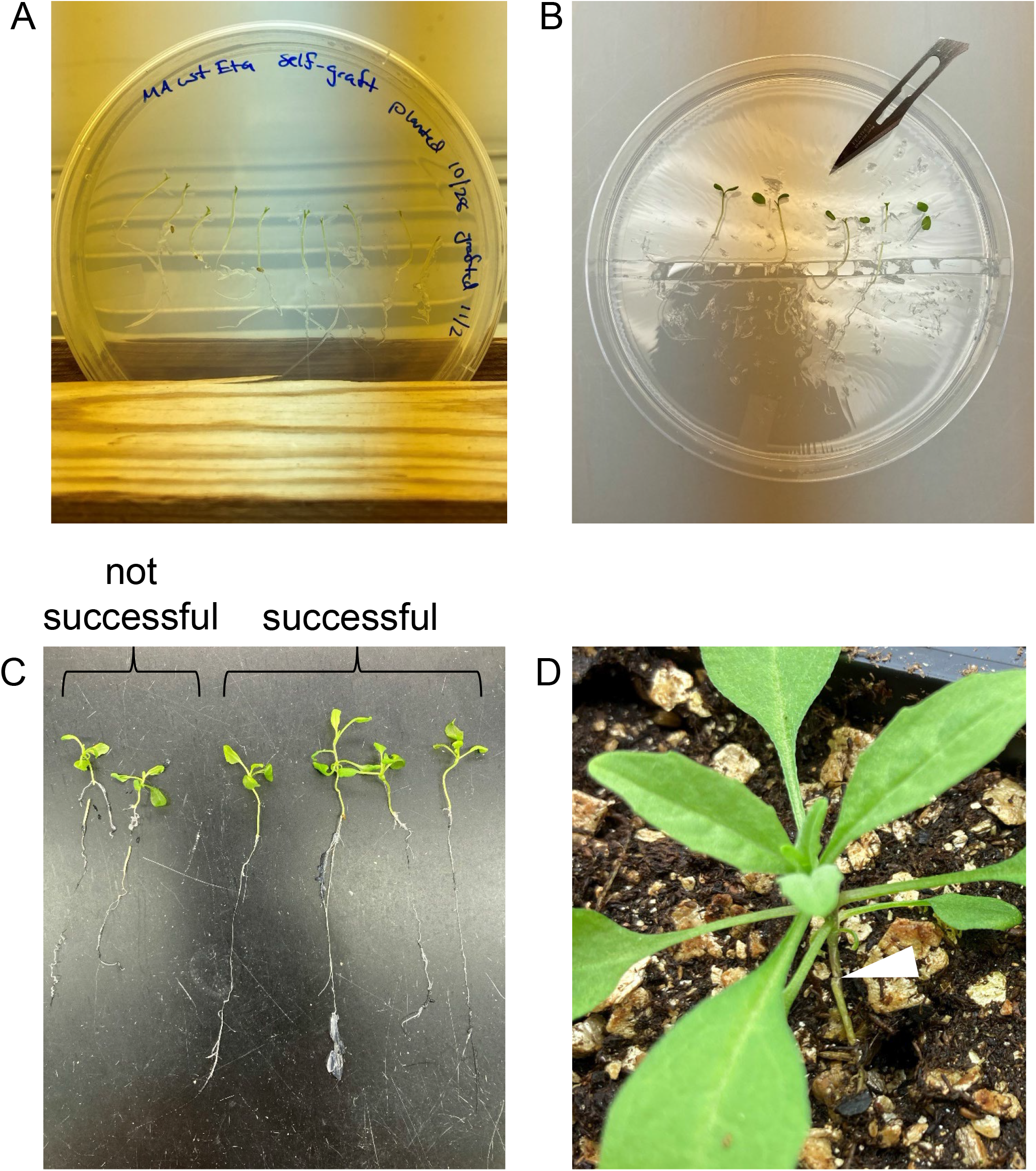
Hypocotyl grafting of *Erysimum cheiranthoides*. (A) The hypocotyls of seedlings germinated on agar are cut. (B) Hypocotyls of stock and scion are placed in contact. (C) In successful grafts, a connection is made between the stock and the scion. In unsuccessful grafts, the scion has made adventitious roots. (D) Successfully grafted plants are transferred to soil. The graft junction is indicated by an arrow. This method is adapted from an Arabidopsis grafting protocol described in Marsch-Martínez *et al*. (2013).

**Supplemental Figure S7:**
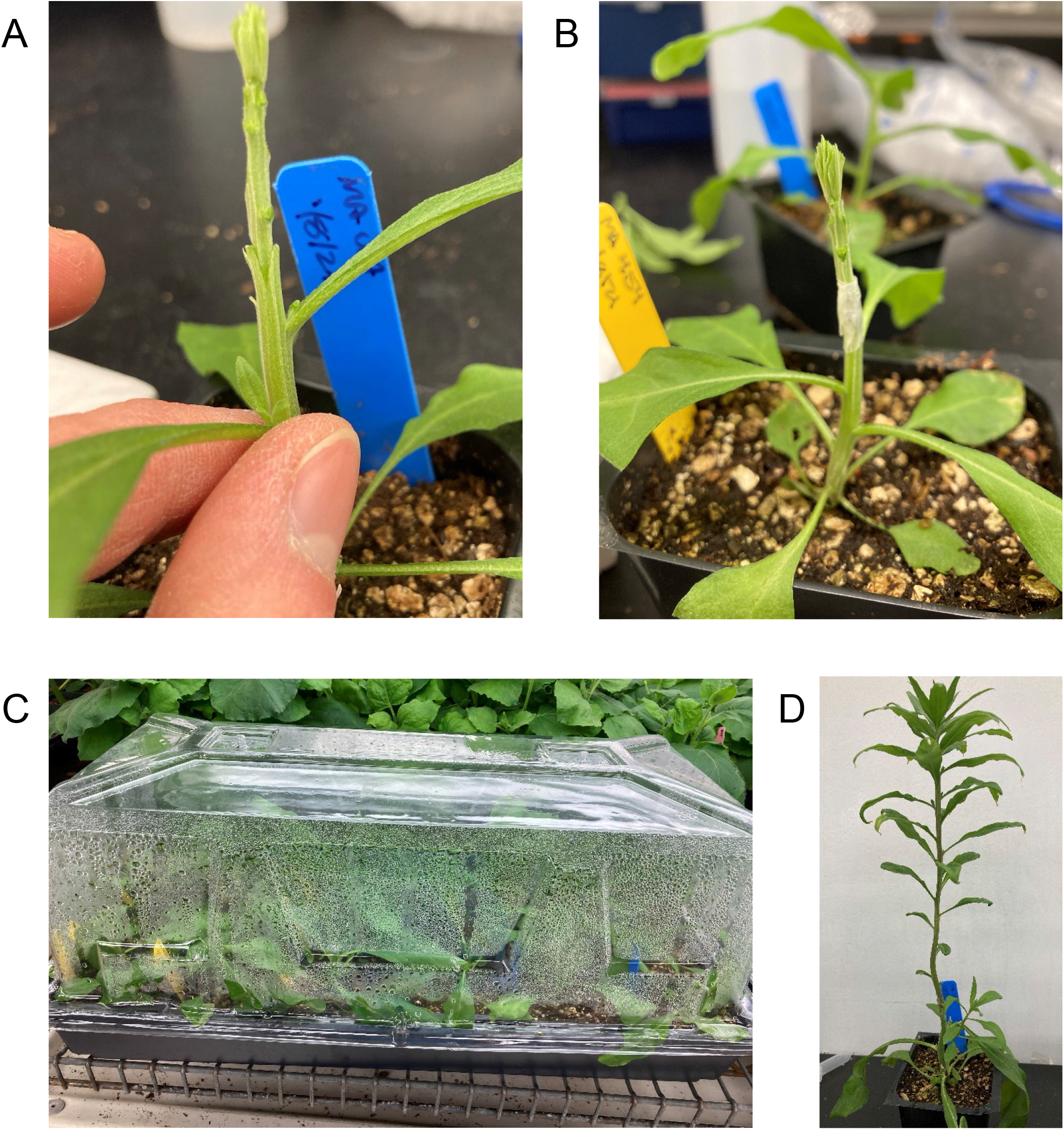
*Erysimum cheiranthoides* juvenile plant grafting. (A) Three to four week-old plants were cut in the middle of the shoot, about four centimeters above the soil line. All expanded leaves were removed from the scion, unexpanded leaves were trimmed to reduce leaf area, and the base of the scion was cut into a wedge shape. One or two leaves below the graft junction were removed from the stock and a vertical cut was made, about 5 mm into the stem. (B) The graft junction was sealed with Parafilm. (C) Plants recovered from grafting under plastic domes to maintain humidity. (D) Recovered plants regain apical growth, and axillary shoots were regularly removed to encourage apical dominance.

**Supplemental Figure S8:**
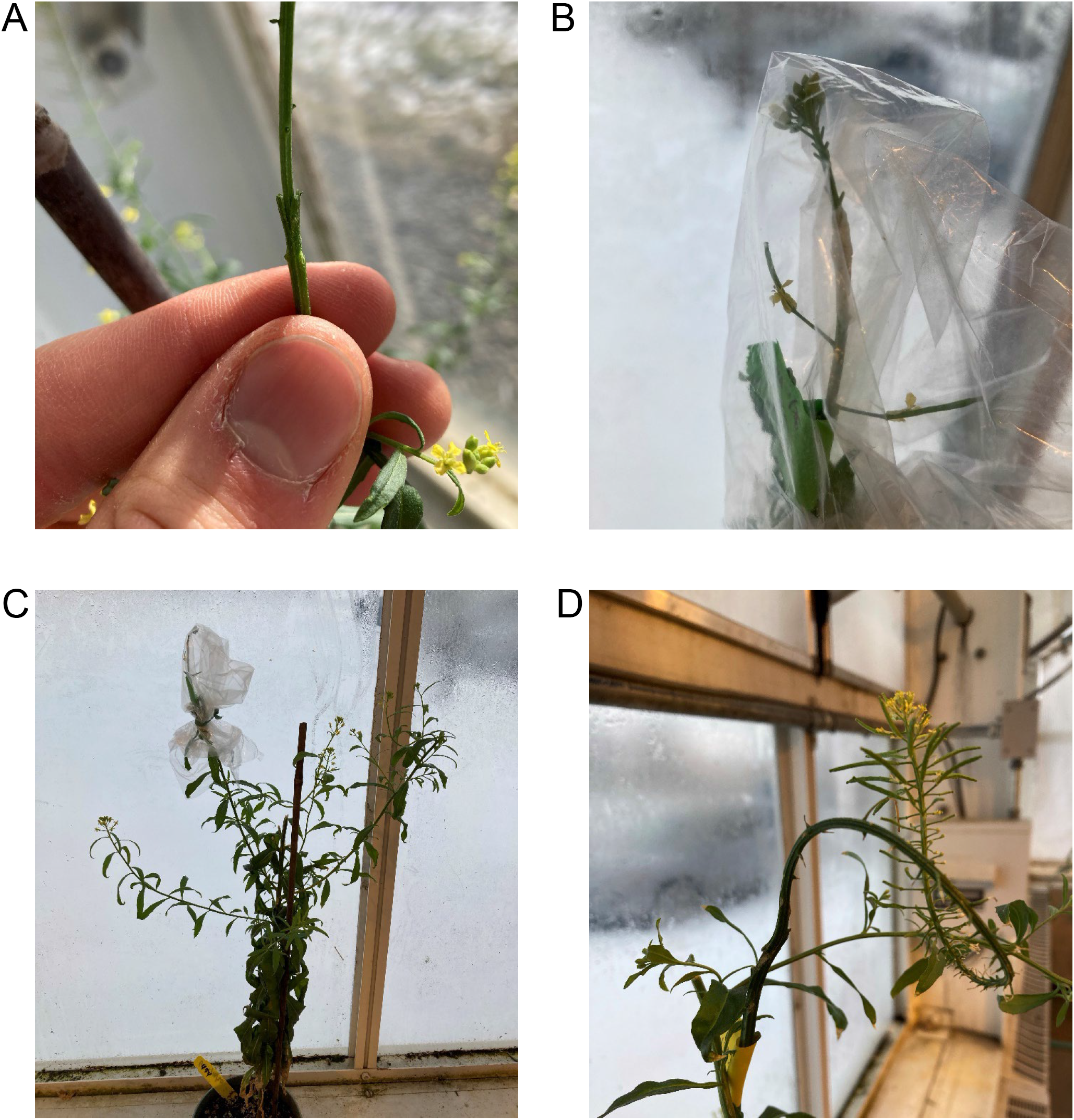
*Erysimum cheiranthoides* flower stalk grafting. (A) Inflorescence apices were cut from *E. cheiranthoides* plants and all siliques, flowers, and buds were removed except for the most distal buds. The base of the inflorescence axis was cut into a wedge shape. The stock inflorescence axis was cut 5 mm vertically and the two axes were joined together. (B) The junction was sealed with Parafilm and the entire inflorescence axis was sealed in a plastic bag to prevent desiccation while recovering. (C) Other inflorescence axes were removed from the plant. Some were left on the plant in this image, but *E. cheiranthoides* regularly produces many more axillary inflorescences. (D) A recovered inflorescence graft. Axis elongation was often slow, as shown in this image, but other grafts elongated similarly to unmodified plants after recovery.

